# Contrastive learning explains the emergence and function of visual category-selective regions

**DOI:** 10.1101/2023.08.04.551888

**Authors:** Jacob S. Prince, George A. Alvarez, Talia Konkle

**Affiliations:** Department of Psychology, Harvard University, USA; Center for Brain Science, Harvard University, USA; Kempner Institute for Biological and Artificial Intelligence, Harvard University, USA

**Keywords:** category selectivity, contrastive learning, functional specialization, population coding

## Abstract

Modular and distributed coding theories of category selectivity along the human ventral visual stream have long existed in tension. Here, we present a reconciling framework – *contrastive coding* – based on a series of analyses relating category selectivity within biological and artificial neural networks. We discover that, in models trained with contrastive self-supervised objectives over a rich natural image diet, category-selective tuning naturally emerges for faces, bodies, scenes, and words. Further, lesions of these model units lead to selective, dissociable recognition deficits, highlighting their distinct functional roles in information processing. Finally, these pre-identified units can predict neural responses in all corresponding face-, scene-, body-, and word-selective regions of human visual cortex, under a highly constrained sparse-positive encoding procedure. The success of this single model indicates that brain-like functional specialization can emerge without category-specific learning pressures, as the system learns to untangle rich image content. Contrastive coding, therefore, provides a unifying account of object category emergence and representation in the human brain.

## Introduction

Our ability to see and recognize objects, people, and the broader environment around us is supported by representations along the ventral visual stream (1–6). Foundational discoveries have charted regions with category-selective tuning, evident for a few categories: faces (7–10), bodies (11, 12), scenes (13–15), and visually presented words (16–18). That is, recorded responses in single units and voxels respond systematically more on average to the neurons’ preferred category, (e.g. images of faces), with only weak responses to other categories. The properties of these regions and the pressures guiding their emergence has been the subject of intense study for decades (19–31). What is the nature of the tuning in these regions that supports the selective responses across images? And, are these category-selective regions better understood distinctly from each other and the representation of other categories, or as interrelated regions that are part of a broader visual representational scheme?

A prominent view of these brain regions is as distinctive, independent, functional modules (32, 3, 19, 33–36). Category-selective regions only exist for a few domains and are thus hypothesized to be different from other visual categories in some important way. For example, for faces, bodies, and scenes: these categories may have particular ecological relevance and may emerge through built-in specialized architectures, supporting domain-specialized tasks such as face individuation (20, 37–39). For the case of word-selectivity, perhaps extensive visual training and a need to interface with long-range connectivity to language regions leads to the emergence of regions such as the visual word-form area (40–42). Brain regions with selectivities for different categories are typically studied in depth by different communities, and are assumed to have very different kinds of visual feature tuning from one another, related purely to their distinct domains (e.g. face-specialized, body-specialized, scene-specialized, letter-specialized). Indeed, some of the strongest evidence for the modularity of these regions comes from selective deficits following brain damage (43–50) or other forms of causal perturbation such as transcranial magnetic stimulation (51–53), electrical microstimulation (54), and pharmacological inactivation (55); see 56 for review). In this way, modular frameworks specify a ‘labeled-line’ type of visual representation (57), where, for example, a neuron’s highly selective firing rates to face images is a direct indicator of its functional role in face processing.

However, modular accounts are restricted to explaining the representation of a few specific visual categories. Empirical evidence shows that not all visual categories have clustered areas of highly selective tuning along the ventral stream (58–60), leaving unspecified how all other categories are represented. Distributed coding accounts are the prominent alternative framework, where the relevant category information is assumed to be encoded in the activation pattern of the large-scale neural population (2, 61–63). Under this view, across the population of neurons, both high and low responses are critical to signal category information (much like how both 1’s and 0’s encode information in a binary code). These accounts sometimes assume the underlying visual feature tuning of units in object-responsive cortex is either fairly generic (e.g. geons or other building-block-like contours that apply to many categories to some degree; 64), or, that the tuning of any one unit may not be particularly interpretable without the context of the full population code (5). This latter perspective is a key element in untangling theories, which tend to de-emphasize the interpretation of the response properties of local units or features, in favor of studying the broader geometry of the population (65). The strongest empirical evidence for a distributed representational account comes from the pervasive category information present across regions of high-level visual cortex (2, 60, 62, 66–68). For example, both face and non-face categories can be decoded from both face-selective regions and non-face selective regions (2, 69). However, it is debated whether this extensive decodable information is all functionally relevant for recognition behavior (70–73).

These modular and distributed representational accounts have continued to develop in parallel, each contributing to a set of empirical puzzle pieces that constrain our understanding of category selectivity along ventral visual cortex. To date, it remains unclear the extent to which these theories are compatible or opposed. For example, one possibility is that there may simply be two modes of representation in the visual system, where a few special categories have more domain-specialized tuning and localist readout schemes, while all others have more generic tuning with accompanying distributed readout schemes. However, further pieces of empirical and computational evidence hint at a deeper unifying relationship between categories with selective regions and the representation of other objects – for example, based on their systematic topographic relationships on the cortical surface (23, 25, 28, 74–76). Along these lines, here we offer an updated account of category-selective regions, which provides specific insight into both the nature of their feature tuning and their function in the context of information readout.

To do so, we leverage a particular kind of deep convolutional neural network model (DNN) to operationalize this unifying perspective. Specifically, we used self-supervised instance-level contrastive learning objectives (77, 78), to train a standard DNN architecture over a large, rich set of natural images (79, 80). The contrastive learning objective does not prioritize any special categories over other object categories, nor does it even presuppose object categories at all. Instead, these self-supervised contrastive models learn to represent every experienced image distinctly from every other image in an embedding space (while being similar to itself under some transformations; see also 81–84). As a consequence, these models develop *emergent* category-level representations: images with similar visual characteristics tend to come from similar categories, and thus are naturally routed through the hierarchy to nearby locations in the representational space.

A key property of contrastive objectives, relevant for the work here, is that the nature of the learned features is intrinsically *diet-dependent*. This notion of “diet” refers to the range of content depicted in the input samples that ultimately govern a model’s learned features. For example, training a contrastive model that experiences only images of faces will learn feature tuning that aims to discriminate among face image content (e.g. 85, 86). Training the model over a richer visual diet, like the ImageNet dataset used here, will provide learned features that aim to discriminate among all 1.2M images (e.g. 81). The set of training images in contrastive models is critical for determining the nature of the learned feature tuning. An important related point is that the set of units within a layer, as a whole, must jointly represent the entire input space. Therefore, the feature tuning of any single unit is influenced not just by the learning diet but also by the tuning of other units in the same layer. In these ways, the feature tuning of each unit in contrastive networks is meaningfully linked to both the scope of visual input and the tuning of other units within and across layers.

Leveraging these contrastive self-supervised models, the aim of the present work is to provide a possible computational explanation of the emergence and function of category-selective tuning, with purely domain-general learning constraints. We show that a contrastive DNN model has emergent category-selective units, which lead to selective and dissociable recognition deficits when lesioned, and which can predict the high-dimensional response structure of diverse category-selective regions in the human brain. We further introduce a sparse positive weighted voxel-encoding scheme, which reflects a more constrained linking procedure between biological and artificial neural network responses, under the hypothesis that the tuning direction of single model units (and neurons) is key for signaling image content. Broadly, we argue that category-selective regions are facets of a rich, diet-dependent, contrastive feature space. To propose a mechanistic account for these signatures, we introduce the concept of positive routing through hierarchical population codes, where units with different tuning serve to channel different content through successive stages into an increasingly untangled latent space.

## Results

### Category-selective tuning emerges in models without category-specialized mechanisms

We first examined whether a contrastive DNN trained without any category-specific architectural motifs or task objectives would show emergent category-selective signatures that mirror those observed in the human ventral visual stream. Note that our use of the term ‘category selectivity’ throughout is specifically referring to the categories (domains) of faces, bodies, scenes, and visual word forms, following the well-characterized category-selective regions of the ventral stream.

We used a popular self-supervised learning objective (Barlow Twins; 77), to train a standard deep convolutional neural network model architecture (AlexNet; 79) using the ImageNet dataset (80). Barlow Twins attempts to minimize the difference in latent space between different augmentations of an image, outputting a high-dimensional embedding of maximally independent dimensions. Although the Barlow Twins algorithm is sometimes described as energy- or covariance-based rather than contrastive (87), it effectively functions as contrastive with respect to encoding dimensions, yielding representations that distinguish between instances (see Methods, 88, 89).

To test for emergent category-selective tuning in the self-supervised model, we designed procedures to mirror the localization of category-selective regions in the human ventral visual stream in typical fMRI experiments. Specifically, we recorded the activations of every model unit in response to the same localizer image set (90) used to identify category-selective regions in the Natural Scenes Dataset (91), and then performed a statistical contrast to identify non-overlapping sets of model units that were selective for faces, bodies, scenes, and words (**Figure 1A**; see *Methods*). This procedure was run separately for every layer of the model, treating pre- and post-ReLU activations as distinct computational stages.

**Figure 1:**
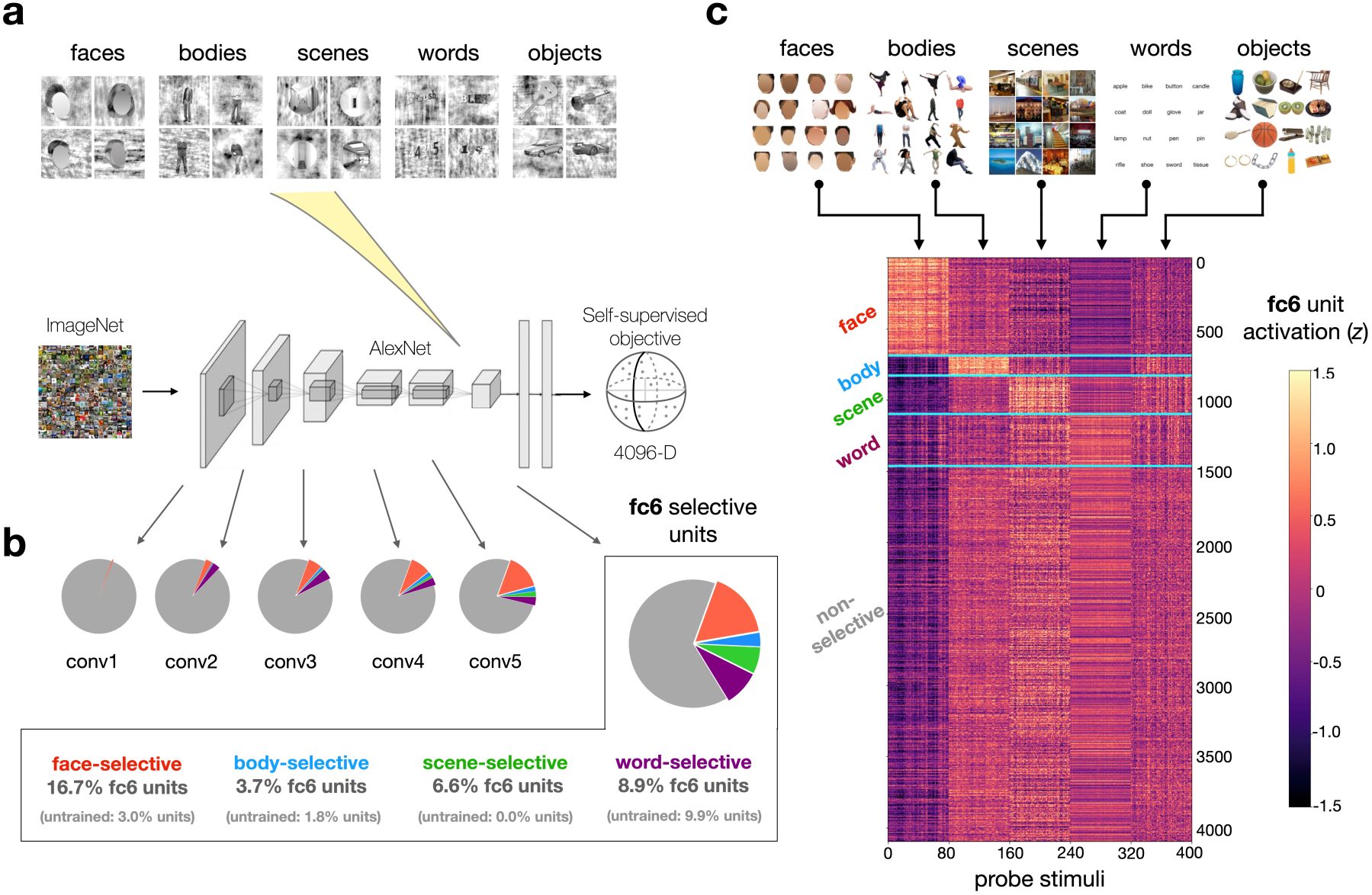
Emergent category selectivity in a self-supervised DNN. (A) Localizer procedure for identifying category-selective units in the self-supervised model. (B) Pie charts summarize proportions of model units selective for faces, bodies, objects, scenes, and words in model layers conv1-5 and fc6 (enlarged). (C) Responses in model layer fc6 to the RGB probe set of face, body, scene, word, and object stimuli. N = 80 images per domain. Layer fc6 units (y axis) are sorted according to their selectivities as defined using the independent localizer stimuli. The 400-dimensional activation profile of each unit is z-scored for purposes of visualization. Images containing identifiable human faces are masked.

With this localizer procedure, we were able to identify small groups of category-selective units in each layer with either robust face-selectivity, place-selectivity, body-selectivity, or word-selectivity (FDR-corrected *p<* 0.05; mean unit *t*-value for faces = 9.29 *±* 4.47 SD; scenes = 8.86 *±* 3.34; bodies = 7.38 *±* 2.17; words = 8.12 *±* 2.74). The overall proportions of selective units tended to significantly increase across the model hierarchy (Spearman *r* = 0.81 between layer index and total proportion of selective units, see **Supplementary Figures 1A, 3A**). The relative strengths of these units’ selectivities also increased significantly as a function of depth (Spearman *r* = 0.92 between layer index and the mean localizer t-value within all selective units; see **Supplementary Figure 2A**). Further, note that even in a single layer (e.g. fc6) it was possible to find different units with selectivity for each domain (**Figure 1B**). Thus, distinct category-selective units emerge in the context of a single population code, as operationalized by a hierarchical layer.

We next tested whether the units’ selectivities generalized to an alternate standard localizer image set. This independent probe set consisted of 400 total images of faces, bodies, scenes, words, and objects, and differed in both low-level image statistics (e.g. color instead of grayscale) and high-level image content (e.g. containing diverse object types instead of only cars and guitars in the ‘object’ category). We observed that the selective units maintained their high responses to preferred stimuli (see **Figure 1C**; **Supplementary Figure 3B**), suggesting that their category-selective tuning is robust and not dependent on a specific probe image set, as is also the case in the human brain. Thus, emergent category-selective tuning is found in a self-supervised DNN model.

Do these signatures of category selectivity depend strongly on the specifics of the localizer method? We tested an alternate procedure, this time identifying units whose mean response to the preferred category is at least 2 times higher than the mean response to each of the non-preferred categories (92). In some layers, this 2:1 approach proved slightly more conservative than the *t*-test method, while in others it was more lenient (**Supplementary Figure 3A**). However, the selective units arising from these two procedures showed highly similar mean activation profiles when probed with independent localizer images (**Supplementary Figure 3B**).

Does category-selective tuning emerge in models trained with other related objectives? Indeed, in an AlexNet model trained with an different contrastive learning objective (Instance-Prototype Contrastive learning—IPCL; 78; see *Methods*), we again observed units across the model with robust selective tuning for each domain, though fewer units selective for bodies (**Supplementary Figure 1B**). Further, an AlexNet model trained using a supervised 1000-way ImageNet classification objective also showed emergent category-selective tuning for the domains of interest (**Supplementary Figure 1C**). Note that the 1000-way supervised objective is also richly contrastive in nature (81, 93)—-its goal is to learn a highly discriminative feature space that can distinguish among many different subordinate categories. However, here we place particular focus on the results of self-supervised contrastive models, for their broader capacity to learn over arbitrary visual inputs without labels, and their added inferential purchase: with self-supervised contrastive models, we can definitively state that no category-level objectives are required for category-selective tuning to emerge.

Finally, since prior reports have indicated that units with face-selective tuning can emerge even in untrained networks (94), we also localized category-selective units in a randomly initialized AlexNet architecture (**Figure 1B, Supplementary Figure 1D**). We found that there were substantially fewer than in trained networks (e.g. 14.6% vs. 35.7% of total layer fc6 units selective for either faces, bodies, scenes, or words in untrained vs. trained models), the strength of these units’ selectivities was far weaker (e.g. fc6 mean *t*-value across domains = 6.6 vs. 13.4, **Supplementary Figure 2A**), and their response properties did not generalize to the independent probe set (**Supplementary Figure 2B**). These results jointly suggest that training (i.e. visual experience) is necessary for DNN features to develop reliable selectivities for these high-level domains of interest.

In summary, the fMRI-inspired localizer procedure was able to identify category-selective DNN units in self-supervised contrastive models. These results demonstrate that units with category-selective tuning to faces, bodies, scenes, and words can emerge as a general consequence of contrastive learning over a rich natural image diet (as reflected in the ImageNet stimulus set). We emphasize that no categorical, semantic or other domain-specific learning pressures were involved in training these models.

### Lesioning category-selective units yields dissociable, predictable deficits

Next we examined the functional role these category-selective DNN units have for image recognition. Neurophysiological evidence has shown that dissociable recognition deficits arise from perturbation of category-selective areas along the ventral visual stream (45, 53, 54), supporting the idea that they reflect distinct functional modules (e.g. 3, 7). If these selective units are also acting as functional modules within the contrastive DNN, then lesioning the units with face selectivity (i.e. setting their activations to 0) should yield a very different profile of recognition deficits than lesioning the units with scene selectivity. Alternatively, these category-selective units may be tuned arbitrarily in the layer’s feature space, and show no clear functional dissociations with respect to the model’s recognition behavior when lesioned.

To explore the impact of lesions of category-selective model units, we first needed to instantiate a readout mechanism to measure the object recognition capacity of the DNN. Note that the self-supervised model is trained only to learn features which discriminate any image view from any other view. How is category information read out from this rich visual feature space? Traditional practices and theoretical frameworks have focused on linear separability of object classes across the full population code (i.e., in the penultimate layer), which is assessed by learning a set of fully-connected readout weights for each category (a distributed readout scheme). However, here, we constrained this procedure by adding sparsity regularization (L1 penalty) to the readout weights. This approach operationalizes a view where category information can be accessed without requiring connections between every neuron in the population and each category node. We offer that this readout method provides a more theoretically constrained and biologically plausible connectivity motif (see *Discussion*). Note that all results below hold with the standard, more flexible fully-connected readout method.

This regularized linear readout function was trained for 10 epochs on top of the penultimate layer embedding (relu7; frozen backbone), and we then measured the baseline recognition accuracy for each of the 1000 ImageNet categories (see **Figure 2A**). Averaging across categories, the mean top-5 classification accuracy was 61.5% *±* 20.3% SD. The learned readout weights were extremely sparse (85.4% of weights with absolute magnitude *<* 0.001, versus 5.4% from unregularized readout with the same hyperparameters), with only a negligible resulting drop in top-5 accuracy of *-*0.95% compared to unregularized readout.

**Figure 2:**
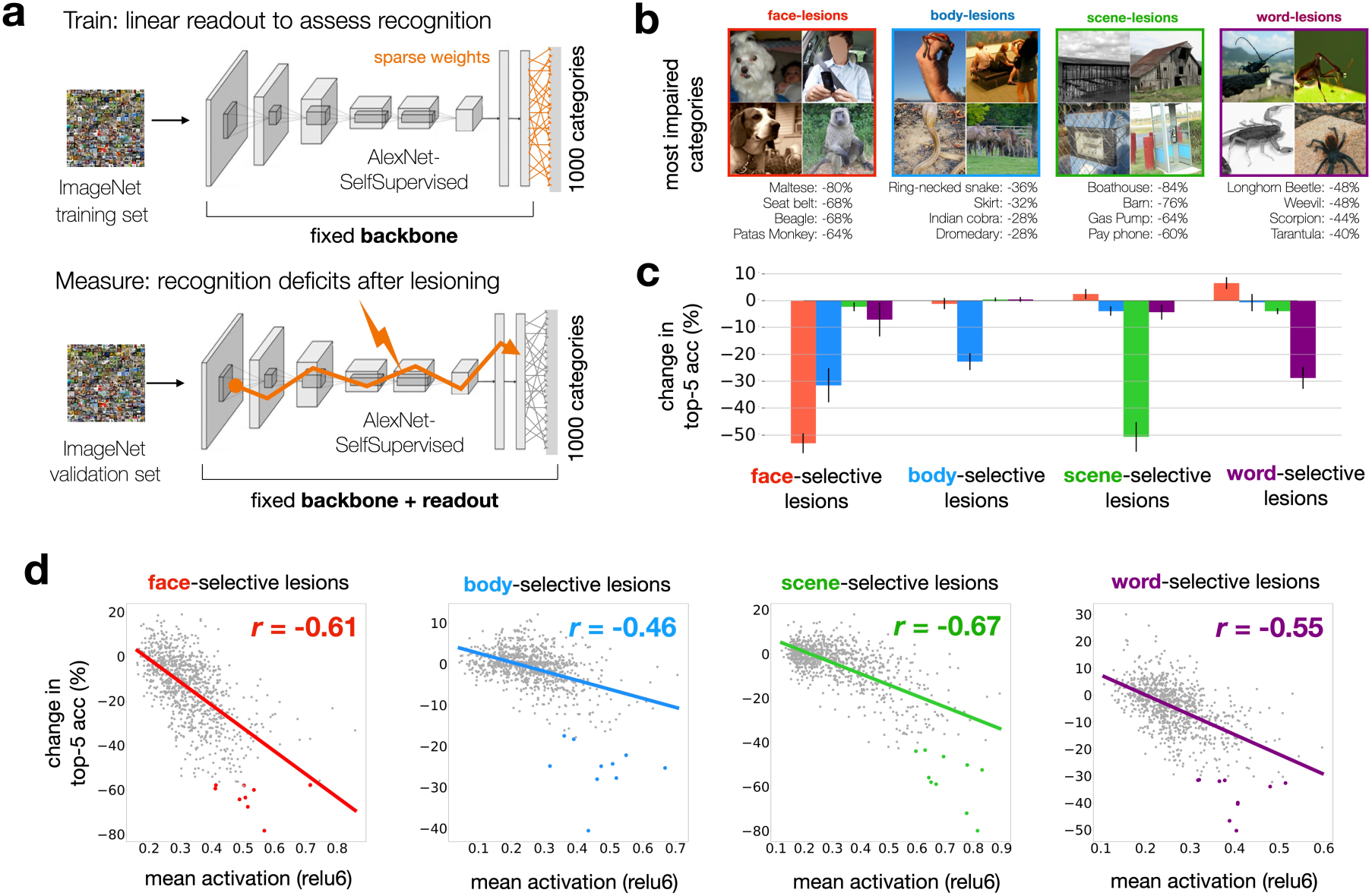
Impact of lesioning category-selective units on object recognition behavior. (A) Procedure for measuring object recognition deficits from lesions to category-selective units in the self-supervised model. Baseline top-5 recognition accuracy is assessed via sparse readout from relu7 features. Recognition deficits reflect the drop in category accuracy following lesions to category-selective units from each domain. (B) Top 4 categories with greatest recognition deficits are shown for lesions to face-, scene-, body-, and word-selective units. Plotted exemplars are chosen from the ImageNet validation set and labeled with drop in top-5 accuracy between baseline and lesioned validation passes. (C) Bar graphs show the mean (± SEM) change in top-5 accuracy over the top 10 categories most impacted by lesions to the selective units of each domain. (D) Relationship between category activation and lesioning cost, for face-, body-, scene-, and word-selective units. Dots reflect the 1000 ImageNet categories. The x axis shows mean activation levels for each category (layer relu6 plotted) from the un-lesioned model. The y axis shows change in top-5 accuracy after lesioning. Colored points reflect the top 10 categories most impacted by lesions to each domain. Y values are jittered (values drawn from normal distribution; mean 0, std 0.5%) to enhance visibility of the results. Photo credit: images in Panel B are samples from the public ImageNet validation set, and those containing human faces are masked.

We next carried out the main lesion experiments. Specifically, we lesioned all of the face-selective units across the model’s relu layers (or body-selective units, or scene-selective units, etc.), and measured the change in recognition accuracy for each ImageNet category after each type of lesion.

To what degree do these distinct lesions produced *dissociable* deficits? To answer this question, we computed the 1000-dimensional cost profiles for each lesion as the difference between unlesioned and lesioned top-5 accuracies for each category. Then we compared these cost-profiles between all pairs of lesions. The cost profiles were broadly unrelated, and in some cases negatively related: as examples, the face vs. scene cost profile correlation was *r* = *-*0.21, face vs. word *r* = *-*0.08, and the average *r* between domain pairs = 0.02 *±* 0.13 SD (see **Supplementary Figure 5B-C**). Thus, it is clear that these different groups of category-selective units have dissociable causal involvement in the recognition of different ImageNet categories.

Which ImageNet categories suffered the greatest recognition deficits for each type of lesion (**Figure 2B**)? It was not obvious *a priori* which categories would be most impacted by each lesion type, especially as ImageNet lacks explicit ‘face,’ ‘body,’ ‘scene,’ and ‘word’ categories, and the ImageNet database was not used in the unit-localization procedure in any way. We observed that lesions to face-selective units led to the strongest deficits for several dog breed categories, as well as object categories containing people, such as ‘seat belt’ and ‘cradle’; body-selective unit lesions led to the strongest deficits for animals and articles of clothing; scene-selective unit lesions led to deficits for certain place categories and large object categories; and word-selective unit lesions led to deficits in recognizing insect categories, as well as objects such as ‘padlock’, ‘wallet’, and ‘dust jacket’ (see **Supplementary Figure 4** for detailed summary of the most impacted categories per lesion). To quantify the strength and selectivity of these deficits, we measured the degree of impairment for the top 10 categories most affected by each lesion type. We used a cross-validation procedure (see *Methods*) to guard against circularity, ensuring that separate images were used to first identify the sensitive categories, and then, to quantify their lesioning deficits. We observed striking multi-way dissociations—for example, the top 10 categories most impacted by lesioning face-units (mean impact: *-*53.2% top-5 accuracy) were broadly unaffected by lesions to scene-units (mean impact: +2.4% top-5 accuracy), with similar trends for the other domains (**Figure 2C**; **Supplementary Figure 5A**). These analyses underscore the distinct functional roles these selective units play within the deep neural network.

The observed impairments bear promising relation to existing findings. For example, fMRI studies have shown that animal images elicit a moderate-to-high response in face-selective regions, and large object categories elicit moderate-to-high responses in scene-selective regions (3, 27, 95). Intriguingly, the occipital word-form area has also been shown to exhibit stronger responses to insects compared to other object classes like chairs or cars (96). The fact that our word-unit lesions highlight insect categories may simply be due to the relative scarcity of letter and word content within ImageNet. However, the link between our results and (96) hints at another possibility: that early experience with thin-lined objects (such as insects) could act as a scaffold for the later formation of letter and word representations. These results set the stage for deeper inquiry into whether causal perturbations of category-selective regions in biological visual system show similar profiles of graded deficits over these images, or, whether they are more specific to a few privileged categories.

Is there a way to predict *a priori* which ImageNet categories are going to be most affected by each lesion? We hypothesized that if a category drives a set of units substantially during normal operation, then its recognition should be substantially impaired when those units are lesioned. That is, we hypothesized that higher activation would be a reliable indicator of greater functional involvement within the network. To test this idea, we examined whether the ImageNet categories that most activated the face-selective units were also the hardest to recognize after lesioning (and similarly for all the other domains). We first calculated the mean activation for each of the 1000 ImageNet categories without lesions, by averaging the activations of the 50 validation set images per category within a given group of selective units. Then, after lesioning those units, we took each category and computed how much its recognition accuracy (again assessed using the validation set) changed due to the lesion. The relationship between these changes in accuracy and the pre-lesion activity levels is plotted for each lesion type in **Figure 2D**. Indeed, we observed consistent negative correlations (mean *r* = *-*0.57 *±* 0.08 SD) – the more an ImageNet category activated a group of category-selective units, the more difficult it was to recognize that category after lesioning. This effect held across layers; see **Supplementary Figure 6A**. The strength and consistency of this relationship implies that positive activation magnitude is a reasonably reliable indicator of functional relevance.

As a key control analysis, we tested the impact of lesioning randomized groups of units. The entire procedure was repeated, this time targeting randomized indices of units that were numerically matched to the size of category-selective subsets in each relu layer. In this regime, we no longer observed meaningful variation in the degree of recognition deficits across categories. As a result, the correlations between activation magnitude and subsequent deficits after lesioning dropped to near zero (**Supplementary Figure 6B**). Thus, systematic deficits for some categories over others only occur when lesioning cohesive sets of similarly-tuned units across the layers.

We also verified that these relationships between activation and recognition deficit were not tied to the specifics of our localizer and lesioning methods. We repeated the analysis using face-selective units chosen via the 2:1-response ratio method, and found equally strong relationships between unit activations and lesion impact (Pearson *r* = *-*0.61 for both localizer methods; **Supplementary Figure 7A-B**). Similar trends were observed when lesioning just the top 1% of most face-selective units from each relu layer (*r* = *-*0.54, **Supplementary Figure 7C**), and when targeting only the face-selective units of layer relu6 (**Supplementary Figure 7D**, *r* = *-*0.60). Finally, to more directly test for a causal link between selective units across the model hierarchy, we silenced only face-selective units across from relu1 through relu5 (leaving subsequent layers unperturbed). Then, we measured the activation levels to the RGB probe localizer images in layer relu6, and observed a strong decrease in activation to face images relative to the other categories (**Supplementary Figure 8A**). The same trends held for the other domains (**Supplementary Figure 8B-D**), suggesting that these similarly-tuned groups of units have a meaningful functional link across the DNN layers. Overall, these analyses show that predictable recognition deficits arise from targeting groups of similarly-tuned units, regardless of their specific locations within the model.

These lesion experiments have two key implications. First, these results demonstrate that functionally dissociable deficits can occur even with a domain-general contrastive objective, and within a general architecture (with no pre-specified modular pathways or branches). Our results imply that during training, the model weights effectively form separable routes through the layer hierarchy, directing different kinds of image content to separable locations in the final embedding space. Second, these analyses reveal a link between a unit’s activation magnitude and its functional relevance in recognition behavior. We hypothesize that this is partially a consequence of the relu nonlinearity, where information propagates from layer to layer through positive activation only.

### Linking model and brain tuning with sparse positive encoding constraints

All experiments conducted so far have focused on a DNN model, identifying units with category-selective tuning, and charting their dissociable functional roles. How similar are the representations in these category-selective model units to the category-selective regions of the brain? We next test the hypothesis that these pre-identified sets of model units from a single contrastive model can capture the response profiles and representational geometries of the classic category selective regions. This departs from many prior approaches, which typically focus on only one category (e.g. only faces, only scenes), often using category-specialized feature models (e.g. models that apply only to faces; 97–100), and, from approaches that train separate end-to-end DNN models to predict each brain ROI independently (30). Instead, here we test the hypothesis that the nature of the feature tuning across all category-selective regions can be understood jointly, within a single model, as a consequence of contrastive feature learning over a rich image diet, in order to distinguish all kinds of visual content.

While most standard approaches model each brain voxel as a linear combination of *all* features from a DNN layer, we instead fit encoding models for each category-selective region using *only* the subsets of DNN units with corresponding selectivity (e.g. only face-selective units are used for modeling FFA and OFA, **Figure 3A**). Further, we impose a second constraint on the encoding procedure, requiring that all learned encoding weights be sparse and positive (**Figure 3B**). Thus, for example, face-selective voxels can only be fit using a minimal positively-weighted combination of face-selective model units. These added forms of regularization strongly enhance the interpretability of the linking function (101).

**Figure 3:**
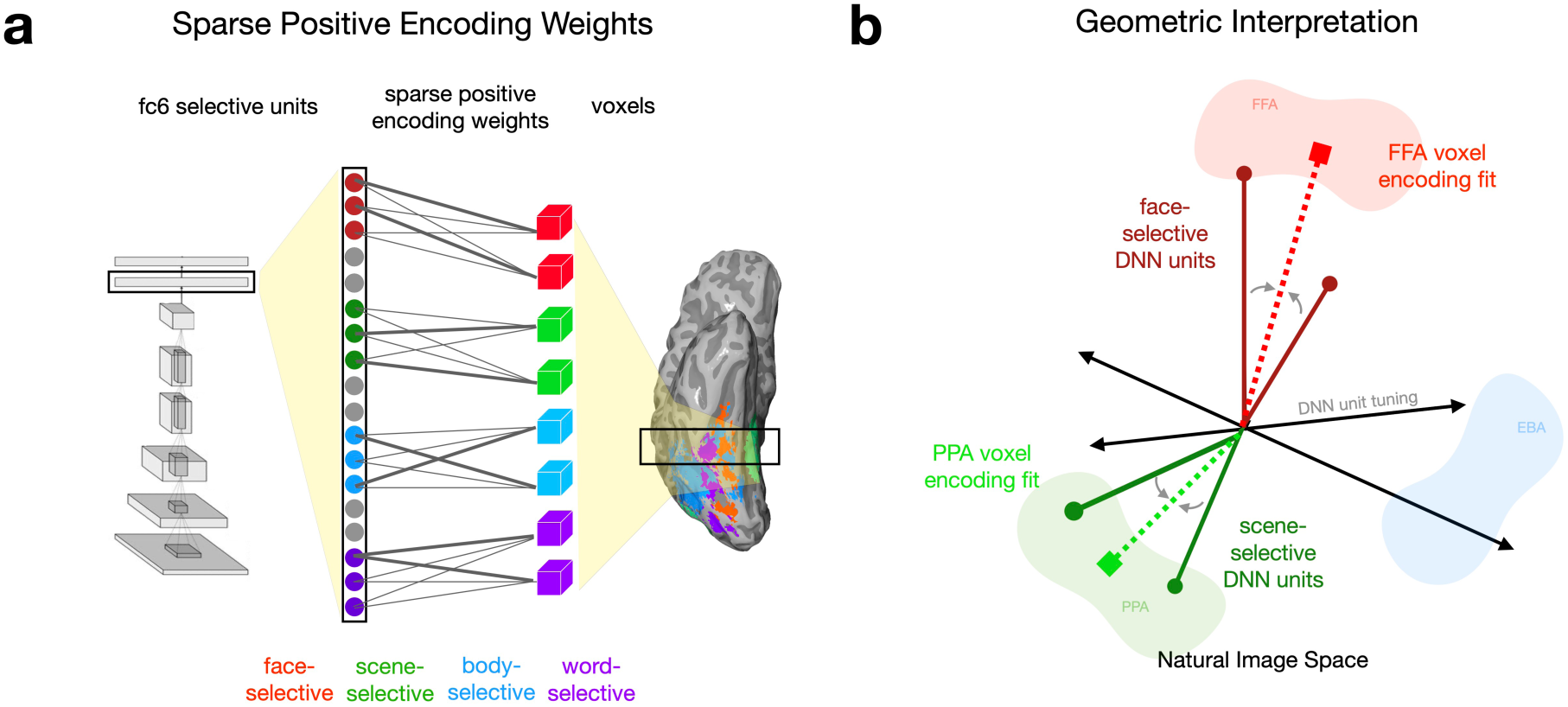
Sparse positive-weighted encoding models of category-selective ROIs. (A) Schematic of the brain-model encoding procedure. DNN units and ROI surface vertices with corresponding selectivity are mapped to one-another via positive-weighted linear regression, and weights are regularized to be sparse (L1 penalty). (B) Impact of the positive weight constraint. In predicting a given brain vertex (e.g. from FFA, shaded red), DNN units with anticorrelated tuning directions to the target (e.g. from scene-selective units, dark green) cannot contribute to encoding fits.

We introduce the sparse positivity constraint on the encoding weights to help ensure that the learned tuning directions between model units and voxels are relatively aligned and in correspondence with each other. *Positivity* is important, as for example, scene-selective and face-selective voxels often have negatively correlated response profiles; standard linking procedures (e.g. linear regression) can apply negative weights to scene-selective DNN units to adequately model face-selective voxels (see **Supplementary Figures 11-12**). *Sparsity* is important to pressure towards as close to a one-to-one alignment between the learned tuning in models and voxels/neurons as possible, and to reduce the amount of feature remixing that happens through the linking procedure. In a way, here we have already implemented a hard sparsity constraint by restricting our voxel-wise encoding models to rely only a small prespecified set of units from the layer. But there are many face-selective voxels and face-selective units, so the additional L1 regularization pushes for an even tighter alignment between voxel tuning and DNN units. Together, these constraints help operationalize the underlying theory of the contrastive account, where the tuning directions of a DNN are meaningfully oriented in the population space, and correspond to the tuning evident across high-level visual cortex.

### Category-selective units account for representational signatures of category-selective ROIs

For our encoding analyses, we used the open-source Natural Scenes Dataset (91), targeting a set of 11 category-selective regions (**Supplementary Figure 9A**). These include 3 face-selective regions (FFA-1, FFA-2, OFA), 3 body-selective regions (EBA, FBA-1, FBA-2), 2 scene-selective regions (PPA, OPA), and 3 visual word-form-selective regions (henceforth, “word-selective”, VWFA-1, VWFA-2, OWFA). All analyses were done on each individual subject. For each voxel in each region, we fit a regularized (sparse positive-weighted) encoding model from the pre-identified model units in each layer with corresponding selectivity, using data from 1000 training images. Using an independent set of 1000 validation images, we then ran a subject-specific cross-validation procedure to identify the most predictive layer (see *Methods*). Finally, we used these voxel-wise encoding models to generate predicted activations to a common test set of 515 images (**Supplementary Figure 9B**).

Our two key outcome measures were the correlation between the predicted and measured univariate response profiles (515 images) and multivariate representational dissimilarity matrices (RDMs, 132,355 pairwise comparisons). Thus, the NSD dataset offers a substantially richer characterization of item-level representational structure in each region than has previously been possible (though there are also clear limitations of this test set, see *General Discussion*).

**Figure 4** shows results for a primary set of 4 ROIs (FFA-1, PPA, EBA, VWFA-1), for both univariate response profiles and multivariate RDMs. We found that the face, scene, body, and word-selective model units each accounted for substantial structure in their corresponding brain regions, for both univariate responses (e.g., 8-subject mean best-layer *r* = 0.58 for FFA-1, *r* = 0.69 for PPA, *r* = 0.48 for EBA, *r* = 0.28 for VWFA-1) as well as multivariate responses (e.g., 8-subject mean best-layer *r* = 0.47 for FFA-1, *r* = 0.46 for PPA, *r* = 0.45 for EBA, *r* = 0.41 for VWFA-1). Here we plot the results from the best-fitting layer, however we note that units from several mid-to-high layers achieved comparable levels of prediction; these results are not highly dependent on the choice of one model layer (see **Supplementary Figures 11-12** for layer-wise results showing similar prediction outcomes across the full set of 11 ROIs; **Supplementary Figure 13** for summary of top-predicting layers). Further, these results hold in parallel analyses conducted in the AlexNet trained on the IPCL objective, as well as on the category-supervised objective (**Supplementary Figure 10**). Thus, the category-selective feature tuning that naturally emerges in a contrastive model is able to well predict the rich and graded brain response structure of all category-selective regions.

**Figure 4:**
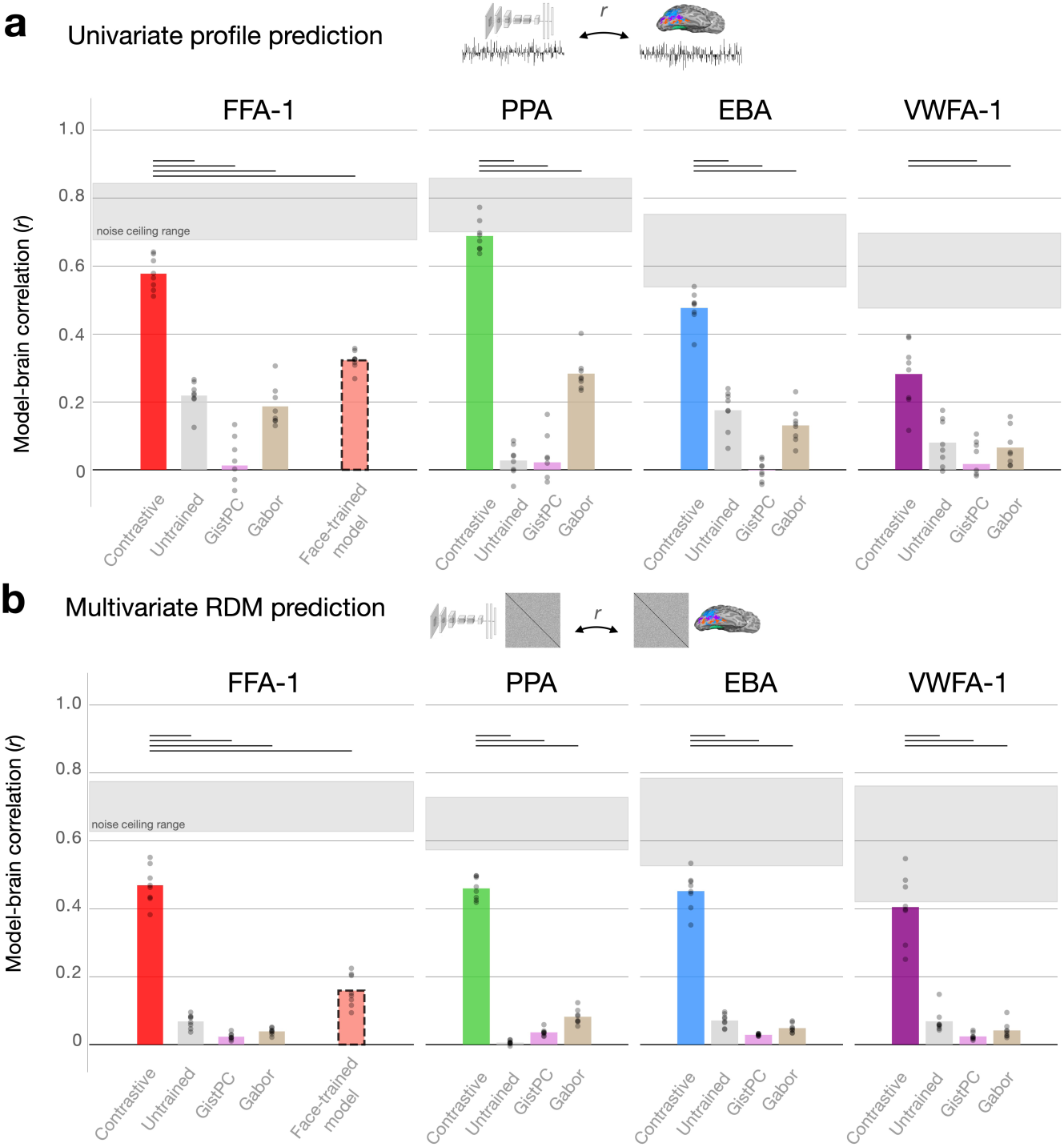
Representational correspondence between model and brain. Encoding results for predicting univariate response profiles (A) and multivariate RDMs (B) are summarized for the 8 NSD subjects, for each DNN and low-level image statistic model. Plotted values reflect best-layer correlations, as defined using cross-validation (see Methods). Shaded regions show the range of subject-specific noise ceilings computed over the data from each ROI. Significance is assessed using paired t-tests over subject-wise prediction levels between the AlexNet Barlow Twins model and each other candidate model. Horizontal bars reflect significant effects favoring the Barlow Twins model; Bonferroni-corrected p< 0.001.

We also observe that the most predictive groups of DNN units are nearly always those with matched selectivity to the neural ROI (e.g. face-selective units predicting FFA, scene-selective units predicting PPA; **Supplementary Figures 11A, 12A**). Critically, this alignment arises only as a consequence of the sparse-positive encoding procedure – when repeating our analyses using unregularized linear regression, all domain-selective subsets are able to achieve reasonably high prediction levels (**Supplementary Figures 11B, 12B**). Thus, sparse-positive regularization is necessary to preserve a meaningful analogy between model and brain tuning directions during the encoding procedure, by preventing the arbitrary reorientation of DNN tuning directions in modeling brain data (101).

Will any feature model be able to predict these brain data? We considered the category-selective units in untrained models, as well as two models of lower-level image statistics (Gist PC and Gabor features, see *Methods*). Across the 11 ROIs, these 3 feature models achieved substantially lower prediction levels than the contrastive DNN unit subsets, for both univariate (paired *t*-test of within-subject differences, 31 of 33 comparisons significant, Bonferroni corrected *p<* 0.001) and multivariate comparisons (29 of 33 tests significant). Indeed, these simpler models accounted for virtually no structure of the brain region RDMs.

Finally, we tested whether a feature space designed specifically for face recognition might also show similar (or even better) emergent fits to face-selective brain regions. We trained an AlexNet model to perform supervised 3,372-way face recognition using the VGGFace2 dataset (102, see *Methods*). The later stages of this model can be thought of as operationalizing a face-specialized feature space, with feature tuning learned solely in the service of within-face discrimination, in order to recognize individuals over changes in viewpoint, lighting, expressions, etc. The trained VGGFace model achieved high accuracy at multi-way face individuation (*⇠*84% top-1 accuracy; linear SVM trained on penultimate layer activations to held-out identities from the test set). We used the sparse positive mapping procedure to fit encoding models from the full activation matrix of each layer to each subject’s face-selective voxels (FFA-1, FFA-2, and OFA).

The face-recognition model features were less predictive than the face-selective subsets of the contrastive model (FFA-1: mean *r* = 0.32 vs 0.58 for univariate prediction, *r* = 0.16 vs 0.47 for RDM prediction; same trend in FFA-2 and OFA; paired *t*-test statistically significant for all comparisons; Bonferroni corrected *p<* 0.0014; **Figure 4A-B**, red). This empirical result is consistent with other work comparing face-trained vs ImageNet-trained models (29, 78, 103, 104). Thus, not just any rich feature space can capture the representational structure of these regions—the face-selective tuning emerging in the contrastive feature space is both distinctive from other feature models and better for matching these brain data.

Overall, these results demonstrate that pre-identified unit subsets from a single DNN model can predict the response structure of 11 diverse category-selective regions, capturing both their univariate tuning over hundreds of natural images, and the multivariate similarity structure over hundreds of thousands of pairwise similarity comparisons.

### Visualizing the emergence of category-selective representation

This set of computational and empirical results offers a unifying account of category-selective regions for faces, bodies, scenes, and words, as distinct facets of an integrated population code. Self-supervised contrastive learning over a rich image diet naturally yields rich discriminative, diet-dependent features. These features are not ‘about’ any one category – the learning is entirely self-supervised and does not presuppose categories. Instead, the set of all features in a layer work together to differentiate all kinds of content. By implication, that model units learn robust face, body, scene, and word selectivity implies that these categories have image statistics that are particularly distinctive in the space of natural image views. Further, this tuning arises gradually across the hierarchical stages (**Supplementary Figure 1**), effectively routing images from their (tangled) pixel-based representation to increasingly distinctive and separable parts of the population code.

To provide a graphical intuition for this progressive untangling of implicit category information, **Figure 5** traces the representational similarity of a small probe set of images from four categories, across several hierarchical processing stages of the contrastive model (conv1, conv3, conv5, fc7; **Figure 5A**; see *Methods*). Each dot reflects an image, where images that are far apart in each multidimensional scaling plot reflects the degree that they evoke more distinct activation patterns across the units in a layer. Early stages show a more ‘tangled’ population code where images from different categories are intermixed, while later stages show clearer emergent separability of these categories.

**Figure 5:**
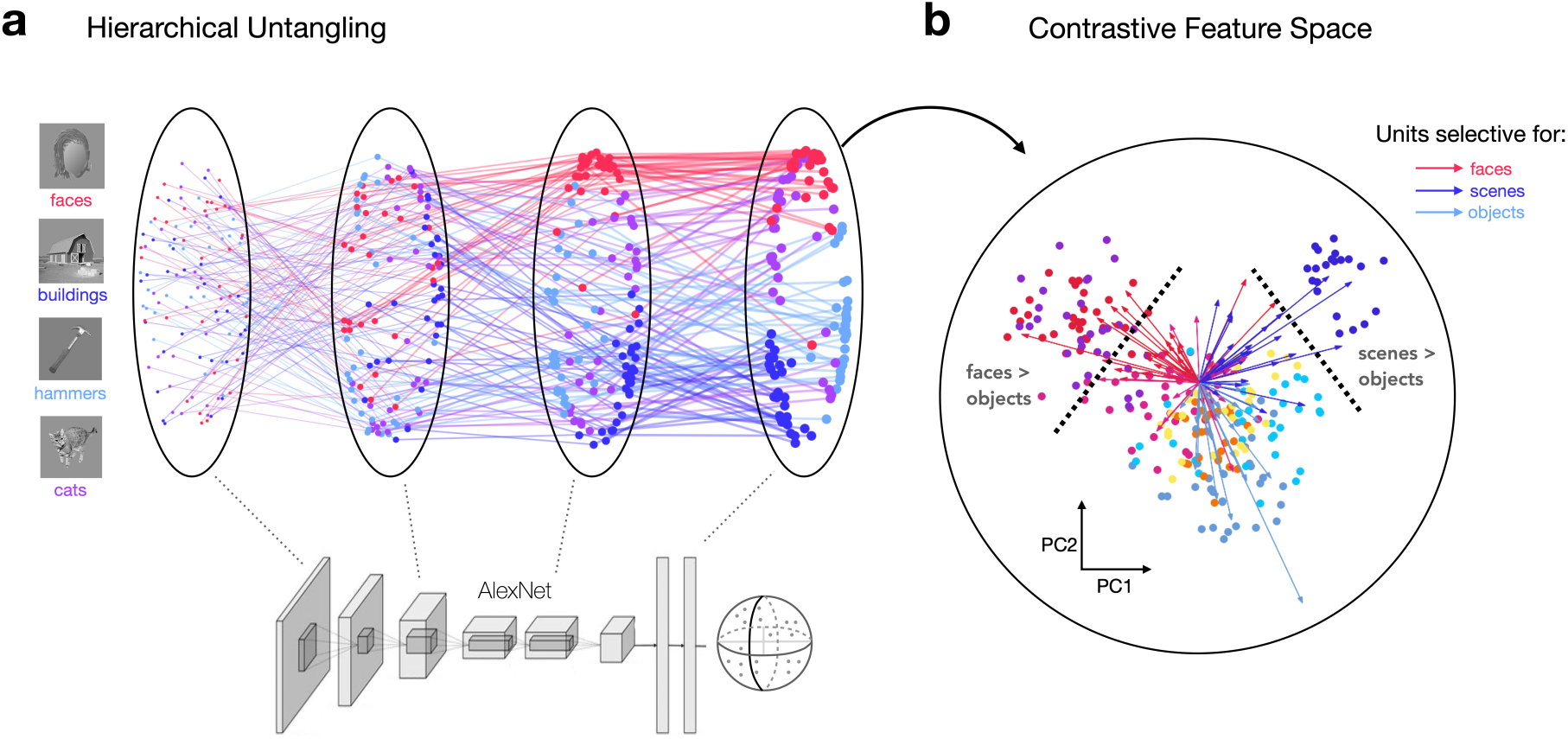
Low-dimensional projection of category-selective axes in a contrastive feature space. (A) Image trajectories through the Barlow Twins model hierarchy are assessed using a probe image set with 30 images from each of 4 categories (105). Each dot reflects the embedding of a single image within a given layer. Only layers conv1, conv3, conv5, and fc7 are included (left to right); each connected sequence of dots shows the hierarchical trajectory of a single image’s feature representations through these 4 layers in MDS space (see Methods). (B) Low-dimensional schematic of category-selective regions in the PC space of layer relu6. Feature representations from the same set of probe stimuli as (A) are plotted in PC2 space; each image is a dot. 4 additional image categories from the same localizer set are plotted: bodies (pink), phones (yellow), cars (orange), and chairs (sky blue). Tuning orientations of the top 25 most selective units for faces, scenes, and bodies are projected into the 2-dimensional PC space of the ImageNet validation set (see Methods). Arrows connect the origin of this relu6 PC space to the 2D projection of each selective unit, with different colors conveying selectivity for faces (red), scenes (dark blue), and objects (light blue-gray). Images containing identifiable human faces are masked.

**Figure 5B** aims to help convey how different category-selective tuning emerges within the same layer. The population code from the fc6 layer is plotted for a probe set of 8 categories (105), along the first two principal components derived from model responses to the ImageNet validation set. In this low-dimensional projection, images of faces, buildings, and bodies naturally have distinctive unit activations from each other as evidenced by the separable clustering. We took the pre-identified face, building, and body-selective model units (from the initial localizer procedure) and projected them into the PC space, indicated by arrows. This visualization helps clarify that units with particular tuning are key for enabling images of faces, bodies, or scenes to separate from all other images in the model’s representational space. Localizing these units with a statistical contrast (e.g. faces *>* objects) is akin to selecting units through a decision boundary in this PC space. These are illustrated with schematic dashed lines in the visualization, highlighting how category-selective units are different facets of this single, unified population code.

## Discussion

The category-selective regions of visual cortex and their relationship to each other and the surrounding occipitotemporal cortex have been the subject of intense debate (e.g. 2, 3). Here we present an updated computational-empirical account of category-selective tuning along the human ventral visual stream. We show that deep neural network models with contrastive objectives, trained over a rich image diet without any labels or category-specific pressures, naturally learn features with emergent category-selective tuning. Further, these emergent model features yield functionally dissociable deficits when lesioned, revealing that implicit modularization naturally arises from very general learning constraints. Finally, the category-selective feature tuning learned across these model units can predict the neural response structure of all corresponding category-selective regions to hundreds of natural images—e.g. predicting face-selective regions better than a model trained solely on face images. Based on these results, we present a unifying contrastive coding account of category selectivity in the brain, arguing that regions like FFA and PPA arise as an emergent consequence of contrastive learning over a rich natural image diet, where positive routing of information along a feature hierarchy implicitly untangles the categorical structure of visual input.

### A contrastive coding alternative to modular and distributed coding frameworks

How are categories represented in the late stage of the ventral visual stream? Modular coding frameworks typically assume that regions tuned to faces, bodies, scenes, and words have distinctive features, specialized for tasks within their specific domains, and that these tuning properties are unrelated to those of other category-selective regions and the surrounding cortex. (32). At the other extreme, distributed coding frameworks typically assume the entire large-scale cortical area is involved in coding for each category, where all regions contribute towards representing all categories through distributed patterns of activity (2). Here, drawing on our computational-empirical results, we present an alternative account where high-level category information is represented with a “contrastive code.” This alternative framework makes distinctive commitments about the (i) nature of the feature tuning in selective regions, and (ii) their functional involvement for reading out category information.

First, what does the contrastive coding framework offer about the nature of the *tuning* in high-level visual cortex? Our account of feature tuning in these regions draws on the principles of contrastive learning objectives, where the learned features are directly determined by the set of visual inputs that must be differentiated. By implication, since the feature tuning is necessarily dependent on the learning diet, the learned selectivities also reflect the scope of the visual input, even in the absence of category supervision. Learning contrastive features from the ImageNet diet necessarily involves learning to distinguish finely among over 1M images depicting a broad range of content. As these contrastive features become progressively more untangled over the hierarchy, they become increasingly selective for specific image distinctions. This allows us to identify units with category-selective tuning using the same category localization procedures applied in classic neuroimaging experiments. For instance, we demonstrate that face-selective units “point to” a specific part of this high-level, contrastive feature space where face images cluster, distinct from the feature directions selective for scenes, bodies, and words. In this way, the precise nature of these contrastive features aligns more with distributed coding theories, since the tuning of any single region fundamentally depends on the tuning across the rest of the population, which collectively works to segregate and distinguish the entire range of visual input.

Second, what does the contrastive coding framework offer about how category information is *read out* of a contrastive feature space? Within distributed coding frameworks, the information is assumed to be spread across the entire population (like a bar code), and thus must be extracted through fully-connected linear readout. In a contrastive space, however, even though the features are part of a shared population code, we show that recognizing individual categories does *not* require fully-connected readout. That is, our implementation of a sparse readout module for category recognition demonstrates that connections to only a small subset of contrastive features can support recognition of any particular category. In this way, we propose that readout from a contrastive code aligns more closely with modular theories, as further demonstrated by the selective and dissociable deficits that arise when lesioning different units. In essence, only a critical subset of units are causally linked to the functional recognition of each category, and these relationships are predictable from high activation levels.

The success of sparse readout from a contrastive feature space has important theoretical implications. This result clarifies that a middle position between modular and distributed frameworks is actually possible, challenging the extreme inference that because there cannot be a neuron for every category in the limit (the grandmother cell hypothesis), the code thus must be fully distributed and read out in a fully-connected manner. Further, from a biological plausibility perspective, sparse readout requires dramatically fewer connections at the interface between a high-level visual feature bank and a cognitive categorical representation. Finally, theoretically, sparse-regularized readout offers an alternative, more constrained, measure of information evident in a population code than the current standard of linear readout (5, 106; see 107, 108 for review).

### Implications of contrastive feature tuning and visual diet

Core to our theory is the idea that the tuning of category-selective neurons in the human brain is fundamentally tied to the statistics of visual experience. We leverage models trained on the ImageNet database, which, while not aiming to be a representative sample of human visual experience, possesses enough scale and diversity for category selectivity for faces, bodies, scenes, and words to emerge. Yet, there are subtle differences between the models and brain data that might stem from the image diet. For instance, human OTC has extensive areas dedicated to body representation (EBA), but our model develops very few units with emergent body selectivity—possibly because the ImageNet visual diet may lack the variety in body pose and social interactions needed to capture the full spectrum of body-relevant features present in human OTC (e.g. 109–114). These points of divergence are now easier to study with the introduction of datasets like EcoSet (115) and SAYCam (infant head-cam data, 116), which reflect more ecologically valid visual experiences, as well as controlled non-ecological image sets (e.g., with all faces blurred, 117). Broadly, this contrastive coding framework lays the groundwork to implement direct manipulations of image diet variation—keeping architecture and contrastive learning objective constant—to identify which aspects of visual experience are critical to account the feature tuning found in high-level visual cortex (118–122, 116, 123). Identifying more ecologically relevant training diets may also support the discovery of new selectivities and content distinctions within occipitotemporal cortex, in a data-informed manner.

In the current study, we test a key visual diet manipulation, comparing the feature tuning in models that receive a diverse ImageNet diet, versus models that only learn to discriminate face identities. We find that the responses of face-selective brain regions to natural images are better captured by models trained on a more varied imageset (see also 29). However, here we have not directly tested whether brain response variation *within* face image sets is also better accounted for by the features of the ImageNet-trained model. On one hand, some evidence suggests this will not hold. For example, in studies of DNN recognition behavior, Dobs et al. have shown that models trained only for supervised object categorization (where faces are one category among others) fail at fine-grained face recognition; specific face-identity training is *required* to accurately classify individual identities (124, see also 125). Given these computational findings, and considering that within-face discrimination is an presumed function of FFA, it is possible that models will require face-specific learning pressures to capture within-face response variation in face-selective brain regions.

On the other hand, we predict that contrastive learning over a diet encompassing both a variety of face identities and object classes should naturally learn features capable of discriminating all these inputs from one another—both differentiating faces from each other and faces from other objects—without needing additional face-specific learning pressures. This remains an open empirical question. Further suggestive evidence comes from studies measuring neural responses to diverse face stimuli in face-selective regions and comparing the predictive capacity of different DNN models. For instance, Vinken et al. examined the responses of face-selective neurons in non-human primates to a large set of face (n = 447) and non-face object stimuli (n = 932), observing that neurons were better accounted for by models trained on ImageNet than by those trained solely on faces (104). Chang et al. observed a similar pattern when studying primate face patch responses to 2100 face images (126). Moreover, in analyzing groups of face-selective neurons recorded intracranially from 33 human patients, Grossman et al. found that ImageNet features more closely matched brain RDMs than VGGFace features for stimulus sets containing only faces (127). Hence, outstanding questions persist regarding (i) whether contrastive learning can achieve individual face recognition without the need for further mechanisms and (ii) whether domain-general contrastive models can accurately fit brain responses of face-selective units when tested with stimuli designed to rigorously probe face identity information.

More generally, this same caveat extends beyond face-selective areas to our broader conclusion that all category-selective regions are part of a unified contrastive feature space, whose feature tuning can be explained without the need for category-specialized pressures. Are the 515 images from the NSD dataset tested here sufficiently representative of natural images to fully support the claim? These images do indeed provide a varied array of natural image content, offering the benefit of not skewing towards a particular representational hypothesis, and reflecting the most extensive probe set of these human category-selective regions to date. However, we fully recognize that more focused tests of the contrastive feature space are warranted, using new or complementary datasets containing many individual face images, body images, or visually presented word images.

### Model-to-brain linking with sparse positive constraints

Our experiments involve two different uses of sparsity: sparse readout (for ImageNet categorization), and sparse-positive linking constraints. These have different motivations. Sparse readout is focused on the internal structure of the model, dictating how one layer interacts with another. This is guided by biological considerations, such as the need for constraints on the number of ‘wires’ or connections required for successful readout. On the other hand, sparse-positive encoding constraints deal with the relationship between two systems – one artificial and one biological. In this case, the objective is to naturally align tuning curves with minimal remixing and reweighting of the input data.

We next discuss the rationale for this constrained *linking* procedure between deep neural network units and brain measurements. Linear encoding models are the current standard approach for relating unit tuning in a deep neural network to voxel or neuron responses. That is, each voxel (or neuron)’s tuning is modeled as a positively and negatively weighted combination of all features in a DNN layer (103, 128–131), or in some cases, across multiple layers (132). Such methods reflect the popular theoretical assumption that the geometry of the feature space is the key determinant of downstream information readout. When considering only the geometry of a space, uniformly rotating all the units’ tuning directions does not affect the information that can be linearly decoded from the population.

However, such rotations would certainly impact which information gets routed from one layer to the next (since only positive activations travel forward in a relu network). Thus, because standard linear encoding procedures allow for arbitrary rotations of tuning directions, they are agnostic to the functional role of tuning (i.e., selectivity) within the broader system (see 133–135 for review).

In contrast, our sparse-positive linking method operationalizes the idea that tuning directions are critical to the function of the system, drawing upon evidence from human neuropsychology (e.g. 56), primate electrophysiology (e.g. 136), and the lesioning results presented here. How does sparse-positive regularization prioritize the role of tuning directions? *Sparsity* (L1 penalty) promotes minimal feature remixing, as it permits only a select few features from the training set to combine and predict the tuning of each target voxel (or neuron). The fact that we restrict encoding of each category-selective region to DNN units with matched selectivity introduces an even stronger sparsity constraint, requiring a direct correspondence between the tuning of selective units in the DNN and selective voxels in the ventral stream. The *positivity* constraint plays a complementary role, limiting the degree to which the sparse encoding weights can rotate or ‘flip’ DNN unit feature directions to map onto brain tuning profiles. Allowing negative encoding weights can lead to counterintuitive and undesirable outcomes, such as scene-selective units achieving high predictivity for (anticorrelated) face-selective ROIs, a phenomenon we document in **Supplementary Figures 11-12**, and in a separate manuscript (101). Together, these sparsity and positivity constraints strongly limit the complexity of the mapping function, providing a more coherent, theoretically informed approach for studying brain alignment.

### A unifying account of visual category selectivity

We have shown that diverse category selective signatures arise within neural networks trained on rich natural image datasets, without the need for explicit category-focused rules. Our findings help reconcile the longstanding tension between modular and distributed theories of category selectivity. Drawing on modular viewpoints, we offer that category-selective regions play a privileged role in processing certain types of visual categories over others. Importantly, we show that domain-specific mechanisms are not necessary to learn these functional subdivisions. Rather, they emerge spontaneously from a domain-general contrastive objective, as the system learns to untangle the visual input. At the same time, our results provide a computational basis for the hallmark observation of distributed coding theories, that pervasive category information is decodable across high-level visual cortex. This widespread decodability is a direct consequence of contrastive learning, since the emergent tuning in one region shapes and structures the tuning in other regions. However, this does not imply that all parts of the population are causally involved in recognizing all visual entities – distributed *representation* does not entail distributed functional *readout*. In recognizing these key distinctions, our contrastive coding framework provides a unifying account of category selectivity, offering a more parsimonious understanding of how visual object information is represented in the human brain.

## Materials and Methods

### Models

Our main analyses involved a standard AlexNet architecture, modified to have group-norm rather than batch-norm operations, which we have found helpful to stabilize self-supervised model training (following 78). The model was trained on the ImageNet dataset using the self-supervised Barlow Twins objective, with the same training procedures and hyperparameters employed in 77. The goal of the Barlow Twins objective is to reduce redundancy in the neural network’s learned representations while preserving the informativeness of its features. In brief, the learning algorithm involves measuring the cross-correlation matrix between the embeddings of two identical networks fed with distorted versions of a batch of image samples, with the objective to make this matrix close to the identity.

Although the Barlow Twins objective has been described as energy- or covariance-based rather than contrastive (87), the objective function emphasizes distinctions across dimensions within a batch, and is thus contrastive with respect to dimensions of encoding. Indeed, the equivalences between these classes of self-supervised algorithms have been extensively validated by studying the properties of their gradients (88) and their generalization behavior (89). For our purposes, the important point is that the Barlow Twins algorithm yields representations that distinguish between instances.

For supplementary analyses with a similar contrastive objective function, the Instance Prototype Contrastive Learning (IPCL) model was applied (78). This self-supervised model also uses an AlexNet architecture but integrates group normalization layers. A forward pass involves creating five augmentations per image, passing them through the DNN backbone, and projecting them into a 128-dimensional L2-normed embedding space. The learning objective includes two components: driving the embeddings of augmented views towards their average representation, and simultaneously encouraging these representations to differ from those of recently encountered items.

For comparison between self-supervised and category-supervised feature spaces, we used the default TorchVision AlexNet model trained with supervised loss on ImageNet. For comparison with a supervised face recognition model, we used another AlexNet variant trained on the VGGFace2 dataset (102). This large-scale dataset contains over 3.3 million images of more than 9,000 individuals, capturing diverse identity variations. The VGGFace trained model was trained using cross-entropy loss and SGD for 100 epochs, adopting the default image transformation scheme.

### DNN localizer procedure

The DNN localizer procedure involved defining category-selective units within each layer of the network by analyzing responses to different categories of images in a localizer image set. Primary analyses relied on the vpnl-fLoc localizer image set (90), which contains 288 stimuli for each of the face, scene, body, object, and character (word) categories, plus 144 additional scrambled stimuli. Different spatial locations in convolutional layers were treated as different units; there was no activation pooling involved in identifying selective units. Face-selective units were identified by comparing activations in response to face images against activations in response to images from each non-face category (bodies, scenes, objects, word, scrambled). Each comparison involved a two-sample *t*-test for each neuron, yielding a *t*-map and corresponding *p*-map reflecting relative preference for faces versus each non-face category. A False Discovery Rate (FDR) threshold was calculated and applied to define ‘selective’ face-preferring neurons for each domain comparison. The intersection of these neuron groups was then identified to define the final set of face-selective units within a layer. This process was repeated for scene-, body-, and word-selective units, ensuring no overlap among groups due to preference requirements.

The robustness of the localizer procedure was evaluated qualitatively by examining responses in layer fc6 to an independent RGB probe set that maintained the same categorical domains (n = 80 stimuli per category), to test for high activations to the preferred stimuli in the corresponding subsets of selective units. To quantify these effects, the generalizability of selectivity estimates was assessed by comparing the *t*-values from the initial localizer procedure with those derived from repeating the same procedure using the RGB probe images. The degree of generalization was reflected in the extent to which the selective units identified using the initial procedure also showed large-magnitude positive *t*-values in response to the RGB probe set. The same generalization analysis was also conducted on an untrained AlexNet Barlow Twins model.

We tested a second method for identifying category selective units, by identifying those with a mean response to the preferred category that is at least 2 times higher than the mean response to each of the non-preferred categories independently. Since this metric derives from fMRI scenarios that compare the (positive) visually-evoked response levels of different localizer categories, here we only compute it within the relu layers of the DNN. We compared this “2:1 ratio” method to the default “t-test + FDR” method by measuring the overlap of selective unit indices by layer (intersection over union).

### Lesioning DNN selective units

This analysis aimed to understand the effect of perturbing groups of DNN units that are selective for each domain (e.g. faces or bodies) on recognition performance. Before introducing any lesions, the model’s baseline recognition accuracy was measured by appending a linear readout layer to the relu7 model layer and training it for 10 epochs on ImageNet categorization. The readout weights were initialized following a normal distribution, with all other trainable parameters frozen. Training was performed using the entire ImageNet training set and employed a batch size of 512. To promote sparsity, an L1 penalty was applied (lambda = 10e-6). A OneCycle learning rate schedule was used, with the following hyperparameters: max lr = 0.05, initial lr = 0.001, pct start = 0.3. After the training phase, recognition accuracy was assessed using the ImageNet validation set, and top-5 accuracy was used for all subsequent analyses.

To quantify the effect of lesions targeting a specific domain, a masking procedure was employed to ‘lesion’ the target selective units in relu layers 1-7, setting their output activations to zero. The indices of the lesioned units for each domain were those identified as category-selective in the DNN localizer procedure. Post-lesioning, the model’s recognition performance was again evaluated, without any further training to assess recovery of category performance. After lesioning, the drop in recognition accuracy levels was evaluated through a second ImageNet validation pass. For our main analyses, all units identified as selective with the t-test + FDR localizer procedure were lesioned. In supplementary analyses, we tested the impact of several alternative lesioning schemes: (i) targeting all units identified as selective with the 2:1 ratio method; (ii) targeting only selective units in layer relu6; and, (iii) targeting only the top 5% or 1% of selective units within relu layers, ranked by their selectivity, rather than the full set of units identified using the localizer procedure.

We sought to identify the *k* categories most affected by lesions to face, body, scene, and word-selective units, and measure how those categories were impacted by lesioning. Critically, ImageNet does not contain explicit “face” or “word” categories, so is no direct correspondence between the localizer-defined units and particular ImageNet classes. Accordingly, we relied on cross-validation to first identify the subset of *k* categories most impacted by each lesion (using half of the ImageNet validation set), and then to independently measure the degree of recognition deficit in each of them (using the held-out half of the validation set). This step helps guard against circularity, by ensuring that our estimates of lesioning deficits are not strongly tied to the particular images used to identify the most-impacted categories. Beyond examining these subsets of categories, we also compared the full 1000-dimensional cost profiles between each pair of domain-targeted lesions, using Pearson similarity.

We tested if activation levels within each group of selective units at baseline could predict subsequent recognition deficits following perturbation. To do so, we computed the correlation between each layer’s 1000-dimensional category activation profile and the lesioning cost profile for each of the face-, scene-, body-, and word-selective lesions. These relationships were assessed for each lesion type, and across our different methods for identifying selective units and implementing lesions.

Finally, we tested the causal impact of lesioning early-layer units on the degree of selectivity observed in higher layers. To do so, we measured the change in activations to the RGB probe localizer set in model layer relu6, after applying lesions to selective units from each domain in only model layers relu1-5. This analysis was repeated twice, once using all units identified as selective in the t-test + FDR procedure, and once using only the top 5% of selective units in each of relu 1-5.

### Relating category selective regions in models and brains

We investigated if category selective units in the DNN could explain the high-dimensional response structure in diverse category selective brain regions. We used the Natural Scenes Dataset (NSD; N = 8; 91), containing measurements of over 70,000 unique stimuli from the Microsoft Common Objects in Context (COCO) dataset (137). We implemented a sparse positive-weighted encoding procedure to map between selective units and brain regions of interest (ROIs) with corresponding selectivities, taking into account both univariate and multivariate response signatures.

For our analysis, we focused on a subset of the dataset (16,515 stimuli; 8 subjects x 2,000 total subject-specific stimuli split equally between the training and validation sets, and a shared 515-stimulus test set). These stimuli, presented during a long-term recognition task, were viewed three times by each subject. We analyzed data from 11 category selective ROIs from occipitotemporal cortex (OTC), defined using an independent localizer procedure. These included face-selective regions (OFA, FFA-1, FFA-2), scene-selective regions (OPA, PPA), body-selective regions (EBA, FBA-1, FBA-2), and word-selective regions (OWFA, VWFA-1, VWFA-2). The same localizer stimuli were used to identify category-selective units in our DNN models.

BOLD responses in NSD were estimated using the GLMsingle toolbox (138). We accounted for potential session-to-session response inconsistencies by *z*-scoring the within-session response profiles in each surface vertex, before extracting responses to the training, validation, and testing stimuli. The data from repeated instances of each stimulus were averaged, and we implemented a reliability-based voxel selection procedure (139) to select vertices with stable response structure. We used noise ceiling signal-to-noise ratio (NCSNR, 91) as our reliability metric and included only voxels with NCSNR *>* 0.3. Surface vertices from the same ROI across the two hemispheres (e.g. lh-PPA and rh-PPA) were concatenated prior to analysis.

Our constrained encoding procedure involved modeling each brain vertex as a linear combination of features from a DNN layer, using only DNN units with corresponding selectivity (e.g. only face-selective units are used for modeling FFA and OFA). All encoding weights were required to be sparse (L1 penalty) and positive. See 101 for further description of this encoding procedure. For the VGGFace model, these constraints only differed in that the entire layer’s features were used for encoding. We conducted an additional set of analyses in the trained Barlow Twins model, comparing matched-selectivity encoding models to those with mismatched selectivities (e.g. mapping scene-, body-, and word-selective units to FFA).

The COCO stimuli were divided into training, validation, and test sets. Features were extracted from each DNN model for each COCO stimulus at each distinct layer of the network from conv3 onward. We used sklearn’s Lasso function to fit each L1-regularized encoding model, enforcing non-negativity of the encoding weights using the ‘positive’ input argument. The default alpha value of 0.1 was used for all encoding fits in the ImageNet and VGGFace-trained models, and a value of 0.001 was used for encoding models from the untrained model.

Once an encoding model was fit, we then predicted the response of every ROI vertex to each image for the validation and test images, and computed two model-predicted outputs for comparison to the true brain responses: the ROI mean univariate response profile and the multivariate similarity structure (“veRSA”, 78, 103). In determining maximal model-brain correspondence, our key metrics were derived from the best-fitting layer to a given brain region. We calculated the univariate and veRSA correlations (Pearson *r*) between model predictions and actual brain data for each layer using the validation set. The layer with the highest correlation was separately selected for univariate and multivariate metrics. Using the independent test set of 515 stimuli, final univariate and veRSA correlations were then computed for the selected layers, providing an independent measure of the maximum correspondence between the model and brain region.

To test if basic image statistic models were able to capture representational structure in the category selective ROIs, we additionally computed encoding fits for Gabor and GistPC feature spaces. Gabor features were extracted in a 16 x 16 grid over the original, recognizable images (425 x 425 pixels) at 4 different scales, with 12, 8, 6, and 4 oriented Gist per scale, respectively (140). This procedure yielded a flattened feature dimension of 7680. GistPC model features were extracted by taking the first 50 principal components of this Gabor feature matrix. The encoding procedures were identical to those described above for the DNN models.

We estimated noise ceilings for each target brain ROI to provide a context for model performance results. We used a recently introduced method based on generative modeling of signal and noise, termed GSN (141), which estimates multivariate Gaussian distributions characterizing the signal and noise structure of each ROI from each subject. Distinct noise ceilings for the univariate and RSA analyses were calculated through Monte Carlo simulations. This process involved first generating a noiseless 515-dimensional univariate response profile (or a 515 x 515 RDM) from the inferred signal distribution. These profiles (or RDMs) were then correlated with counterparts constructed from noisy measurements. The latter were generated by adding samples from the estimated signal and noise distributions, thereby effectively simulating the realistic observational conditions. The range of estimated noise ceilings for the 8 subjects are plotted separately for each outcome metric and each brain region.

### Low-dimensional visualization of hierarchical contrastive feature spaces

To map the hierarchical evolution of category-selective tuning, we visualized the gradual emergence of categorical structure in the Barlow Twins model using a compact image set of four object categories: faces, buildings, bodies, and cats (105). Each category contained 30 examples, with equalized overall luminance and contrast levels across the 120 images, achieved via the SHINE toolbox.

For our hierarchical visualization, we focused on model layers conv1, conv3, conv5, and fc7. We initially condensed the activation data for each layer (120 images x n features) via Principal Component Analysis (PCA) into a 10-component matrix (120 images x 10 PCs). These were subsequently merged into one matrix (480 images x 10 PCs), enabling the analysis of each image’s ‘representational trajectory’ through the model (142). Using Pearson dissimilarity, we then constructed a ‘meta-RDM’ to encapsulate the pairwise similarity of all image features across all layers (480 images x 480 images). The meta-RDM served as input to multidimensional scaling (MDS), yielding a 2D projection of the entire representational geometry (480 images x 2 MDS dimensions).

This 2D representation was then lifted into a 3D visualization. Here, embeddings from disparate layers were grouped into different x-axis positions, symbolizing the progression from conv1 to fc7. The result was a 3D scatter plot where each dot signifies an input image, connected by lines to indicate its trajectory through the model layers. The proximity between dots within each layer’s MDS space reflects the similarity of their corresponding embeddings in DNN feature space.

To demonstrate the demarcation of category-selective regions within this feature space, we created another low-dimensional visualization using the 120-image probe set. We first performed 2D PCA on 4096-dimensional layer fc6 activations computed for a subset of the ImageNet validation set (2000 total images; 2 per category), and then projected the fc6 response vectors for each of the 120 probe stimuli into this 2D space, with dots color-coded by category.

Lastly, we aimed to visualize the tuning directions of category-selective units within this low-dimensional projection. Using the output of the DNN localizer procedure (vpnl-fLoc image set, 90), we identified the indices of face-, body-, and scene-selective units in layer fc6 of the Barlow Twins model. We then depicted the oriented tuning vector of each of the 25 most selective units for these three domains. This was accomplished by creating a one-hot 4096-dimensional vector denoting the index of the unit, and computing its dot product with the (4096, 2)-dimensional PCA component matrix from the ImageNet validation stimuli. A constant scaling factor of 2.5e4 was applied to each one-hot vector before multiplication for visualization purposes. This ensured the resultant tuning vector’s magnitude laid meaningfully onto the image data points in PC space, without altering the relative angles of the different tuning vectors.

## Acknowledgments

We thank all members of Harvard Vision Lab, K. Kay, K. Vinken, G. Tuckute, and N. Blauch for helpful conversations throughout the project and for feedback on the manuscript.

## Author Contributions

Conceptualization: JSP and TK. Methodology: JSP, TK, and GAA. Investigation: JSP, TK, and GAA. Visualization: JSP and TK. Supervision: TK and GAA.

Writing—original draft: JSP and TK. Writing—review and editing: JSP, TK, and GAA.

## Data and materials availability

Public image datasets used to train the models include: ImageNet (https://image-net.org/) and VGGFace2 (https://github.com/ox-vgg/vgg face2). All brain data analyzed here from the Natural Scenes Dataset are available here: https://naturalscenesdataset.org/. All protocols and resources needed to evaluate the conclusions in the paper are present in the paper, the Dataverse repository https://dataverse.harvard.edu/dataset.xhtml?persistentId=doi:10.7910/DVN/5848EQ, and on GitHub: https://github.com/jacob-prince/PROJECT DNFFA and https://github.com/jacob-prince/jsputils.

## Supplementary Materials

**Supplementary Figure 1:**
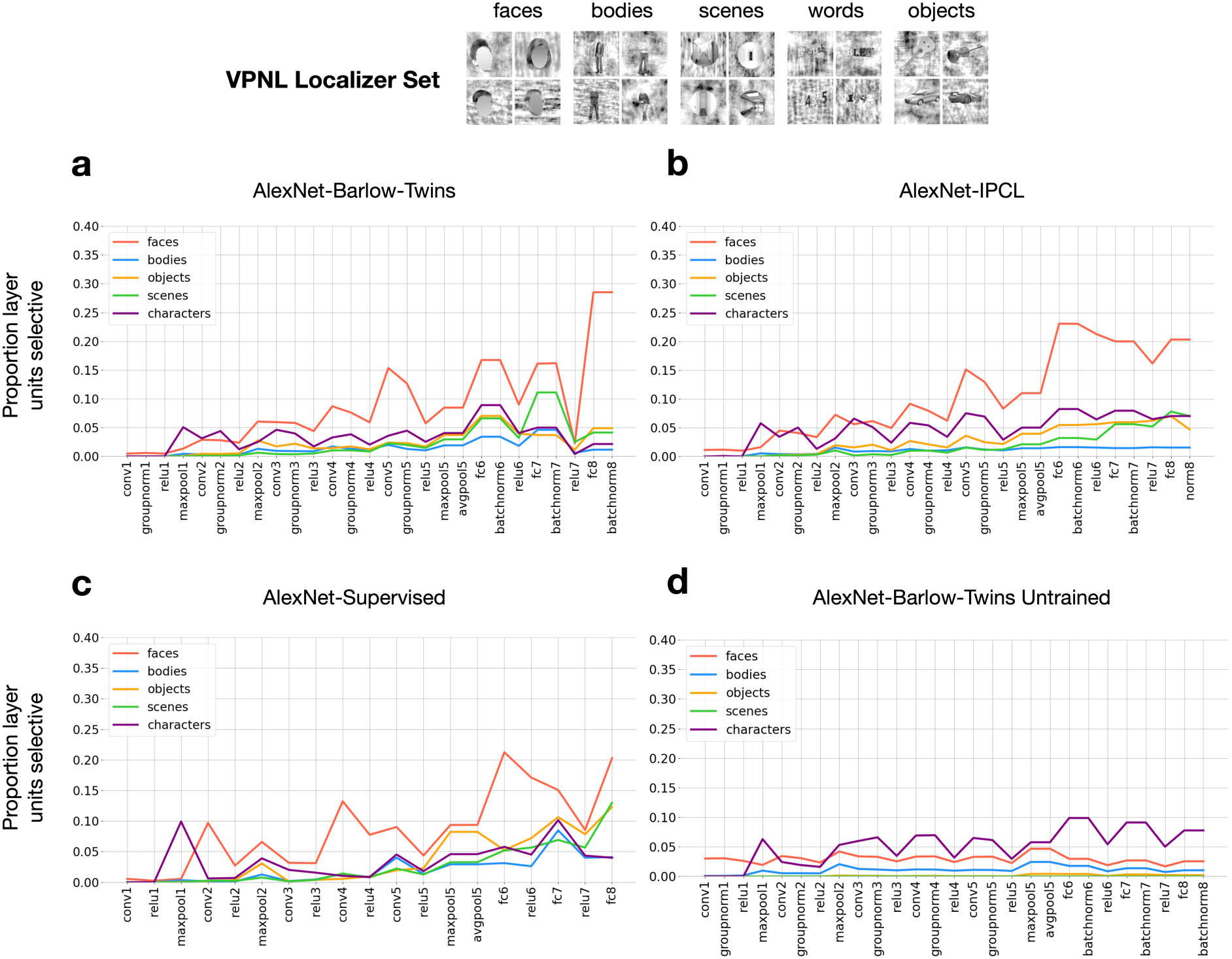
Layer summary of localizer outcomes across DNNs. Layerwise proportions of units selective for the 5 localizer domains are plotted for the trained AlexNet Barlow Twins model (top left), the AlexNet IPCL model (top right), a category-supervised AlexNet ImageNet model (TorchVision pretrained implementation; bottom left), and an untrained initialization of the Barlow Twins model (bottom right). Images containing identifiable human faces are masked.

**Supplementary Figure 2:**
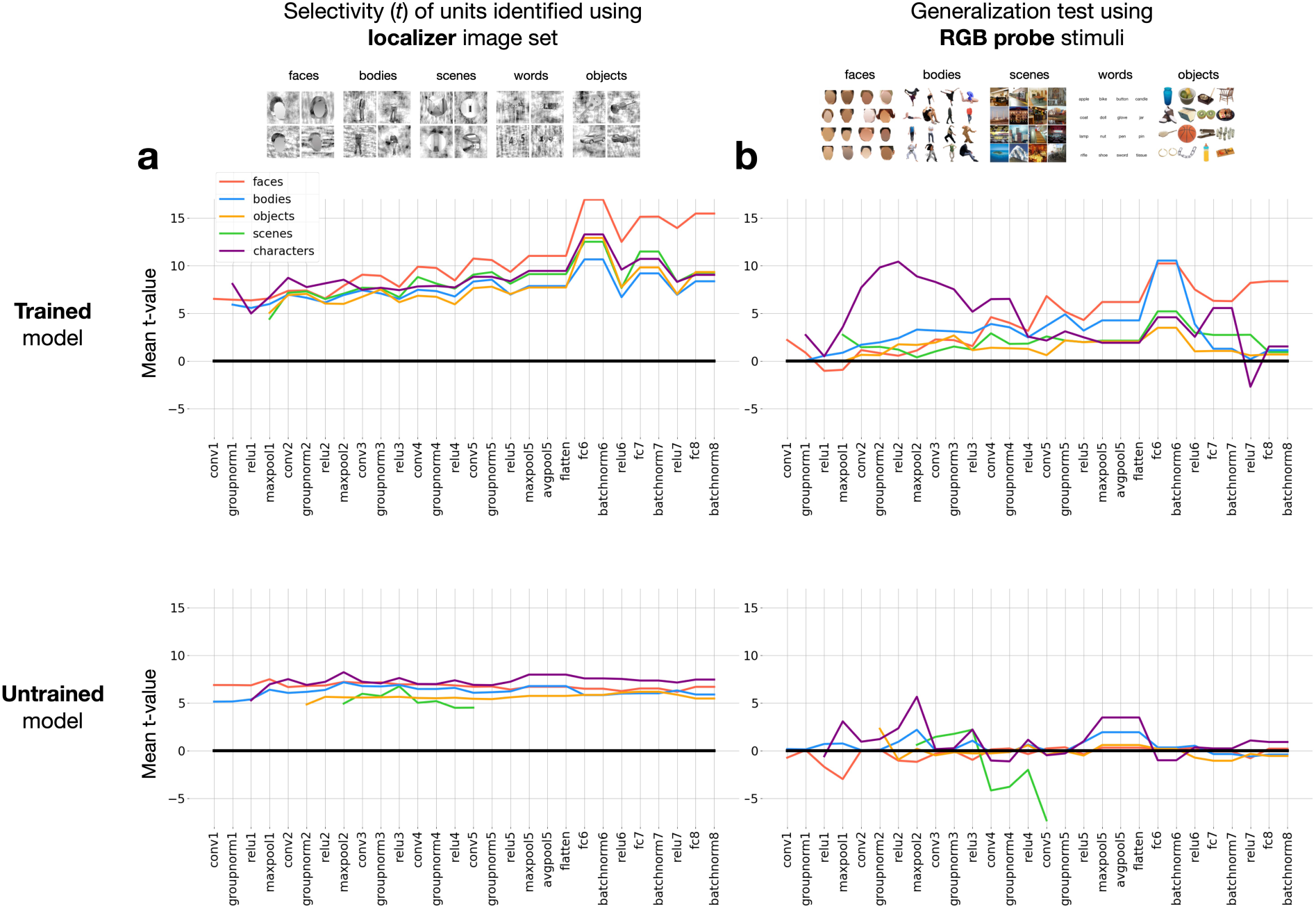
Selectivity profiles do not generalize in an untrained DNN. (A) Strength of units’ selectivities for their preferred domains from the initial localizer run (vpnl-fLoc stimuli). Selectivities are summarized across model layers for the trained AlexNet Barlow Twins model (top) and the untrained model (bottom). The y-axis shows the mean t-statistic within each group of selective units (averaged both over units and over all statistical contrasts involving the target domain, e.g. faces vs. scenes, faces vs. bodies, etc.) (B) Generalization of selectivity to the independent RGB probe set. The t-values are reported from the same groups of units (defined using the vpnl-floc set), with the contrast t-values now computed over activations from the RGB image set. Images containing identifiable human faces are masked.

**Supplementary Figure 3:**
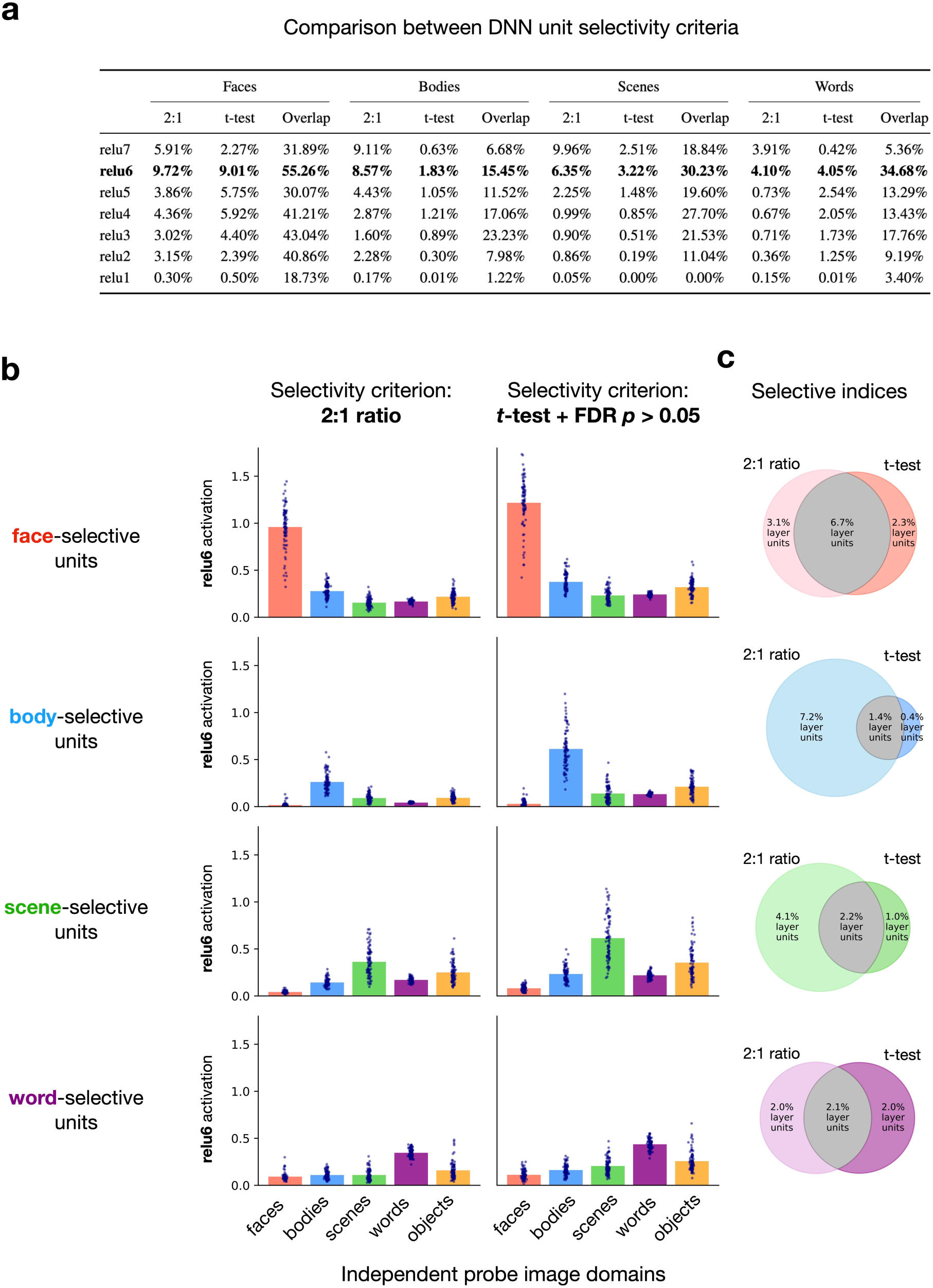
Comparison of selectivity criteria. (A) Outcomes of the 2:1 ratio and the t-test + FDR methods for defining selectivity. Proportions of selective units are shown for all relu layers of the AlexNet Barlow Twins model. ’Overlap’ refers to the intersection over union of unit indices. (B) Comparing selective unit activations to RGB probe localizer images in layer relu6, between selectivity criteria. Bars reflect the mean activation within each group of units over the 80 images from each localizer domain. Image-wise means are plotted as dots. (C) Overlap of selective unit indices between the selectivity criteria, shown as venn diagrams.

**Supplementary Figure 4:**
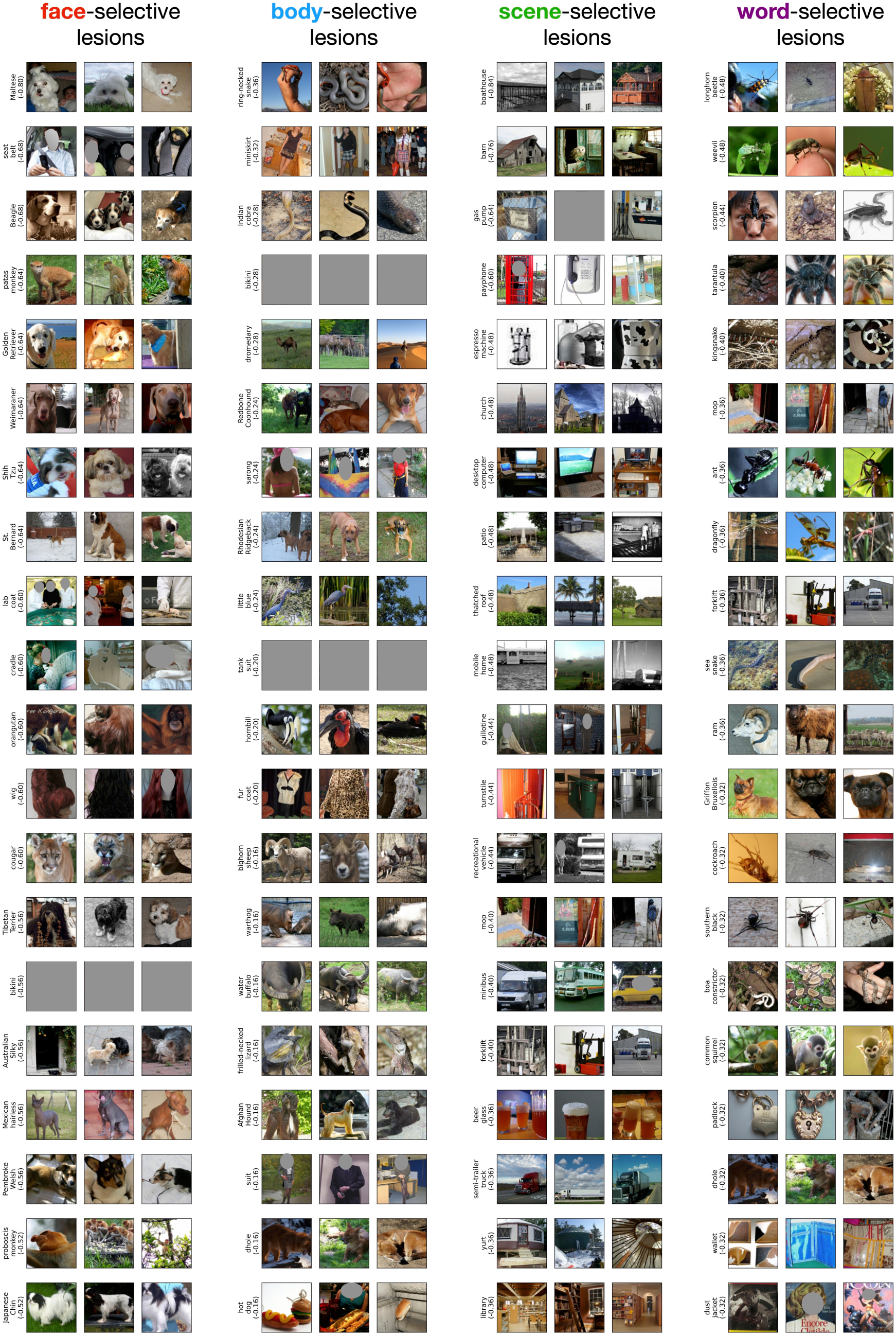
Categories with recognition most impaired by each lesion. The top 20 categories most impaired by each lesion type are identified using cross-validation (see Methods). Photo credit: images are samples from the public ImageNet validation set, and those containing human faces are masked. Some images are fully obscured due to their content.

**Supplementary Figure 5:**
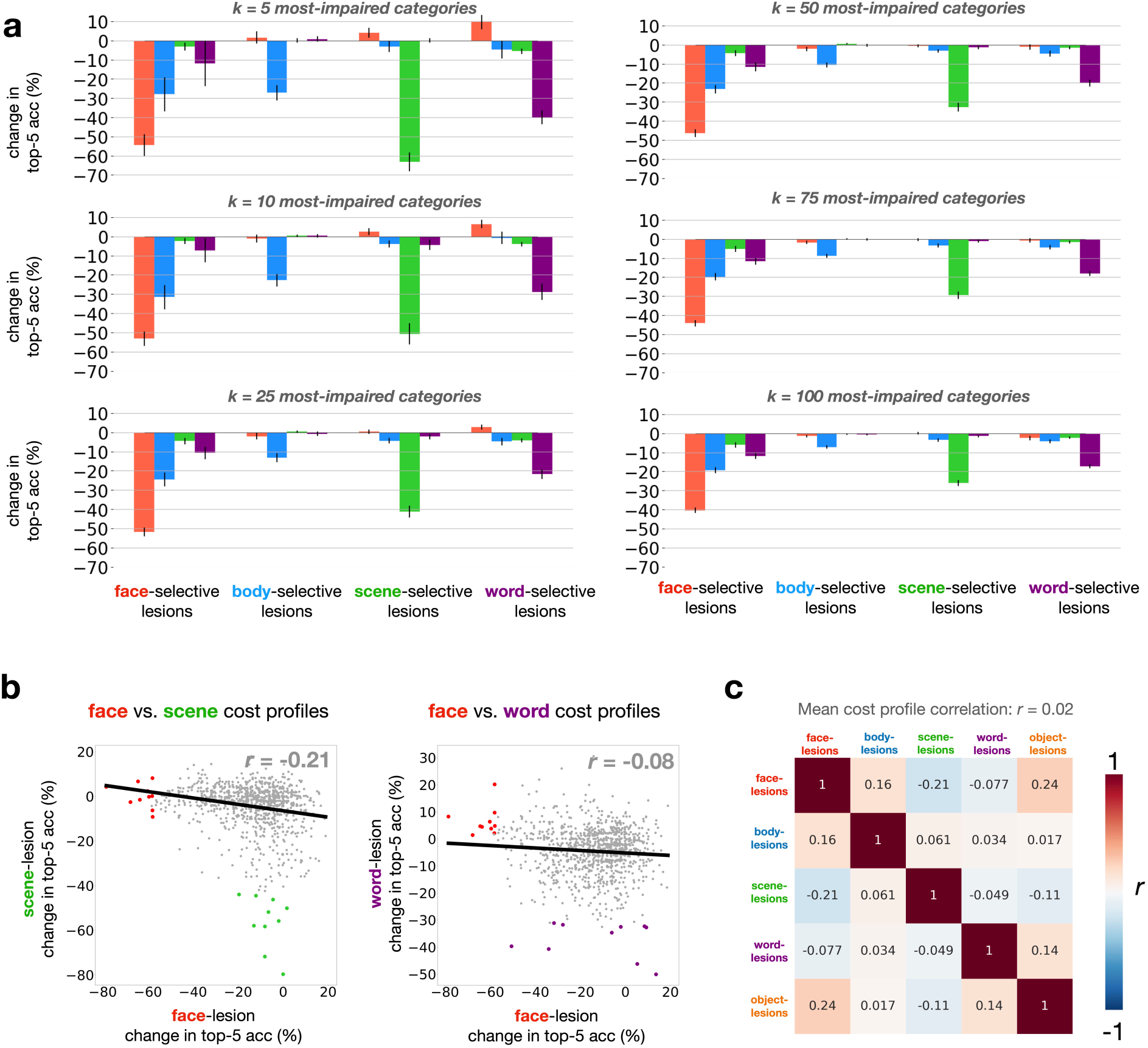
Summary of dissociable lesioning deficits. (A) Bar graphs show the mean (± SEM) change in % top-5 recognition accuracy over the top k categories most impacted by lesions to the selective units of each domain. Indices of the k categories are identified using half the ImageNet validation set, and accuracies for plotting are computed using the other half. (B) Relationship between 1000-dimensional profile of category-level lesioning deficits for face vs. scene unit lesions (left) and face vs. word unit lesions (right). Colored dots reflect the top 10 categories impacted by each domain’s lesion. X and Y values are jittered (values drawn from normal distribution; mean 0, std 0.5%) to enhance visibility of the results. (C) Full pairwise similarity matrix (r) comparing the 1000-dim profiles of lesioning deficits that arise from lesions to each domain of selective units.

**Supplementary Figure 6:**
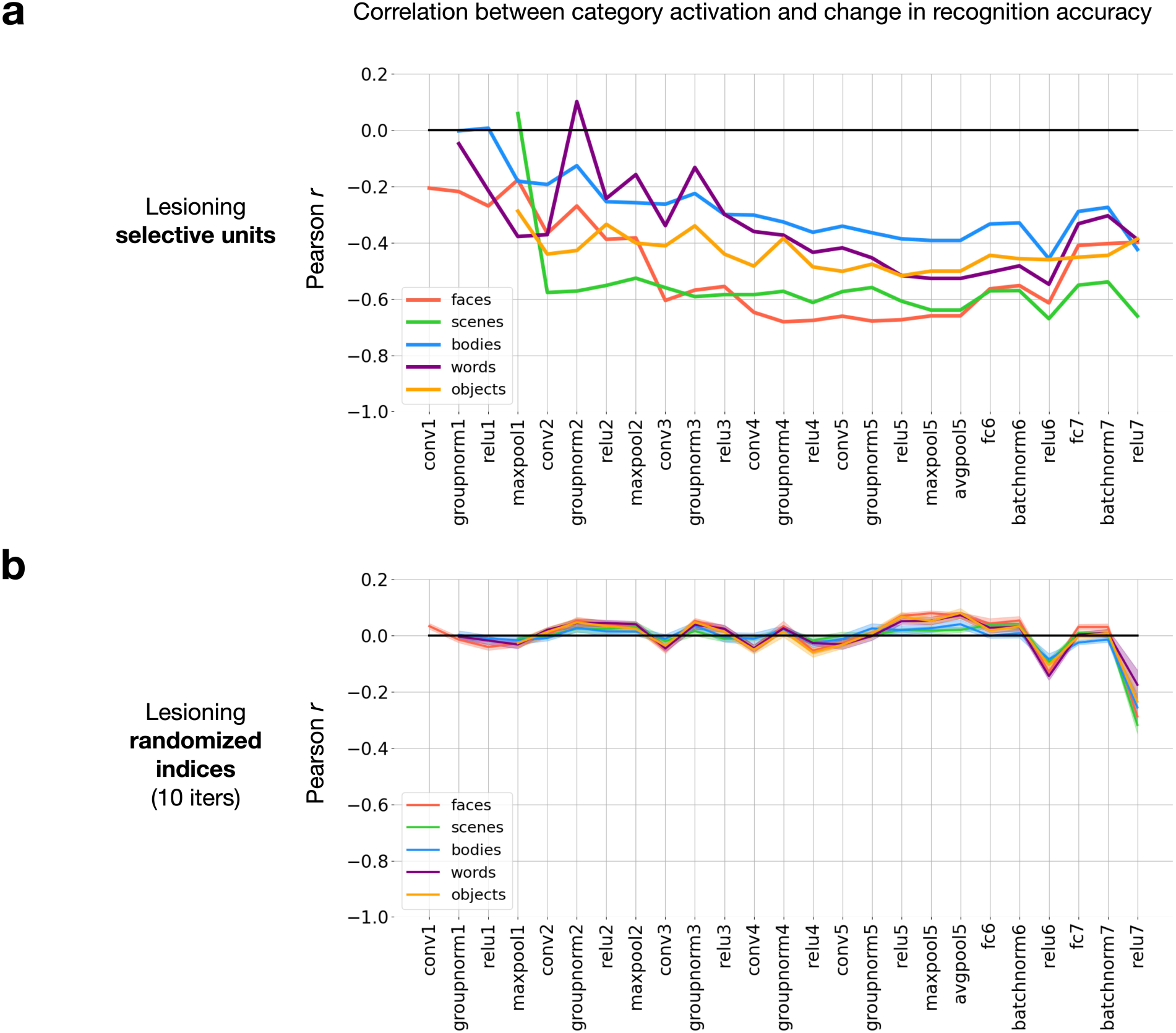
Relationship between activation and recognition deficit for selective-unit and randomized lesions. (A) Layer summary of the relationship between 1000-category profiles of mean activation magnitude (in the unlesioned model) and the changes in recognition accuracy observed after lesioning each domain’s selective units. Data plotted are from the AlexNet Barlow Twins model, and lesions are applied to layers relu1-7. (B) The same analysis as (A), except that the indices of target units are randomized (within each layer) prior to lesioning. The randomized lesion experiment is repeated 10 times, with the mean ± SEM of resultant correlation values plotted.

**Supplementary Figure 7:**
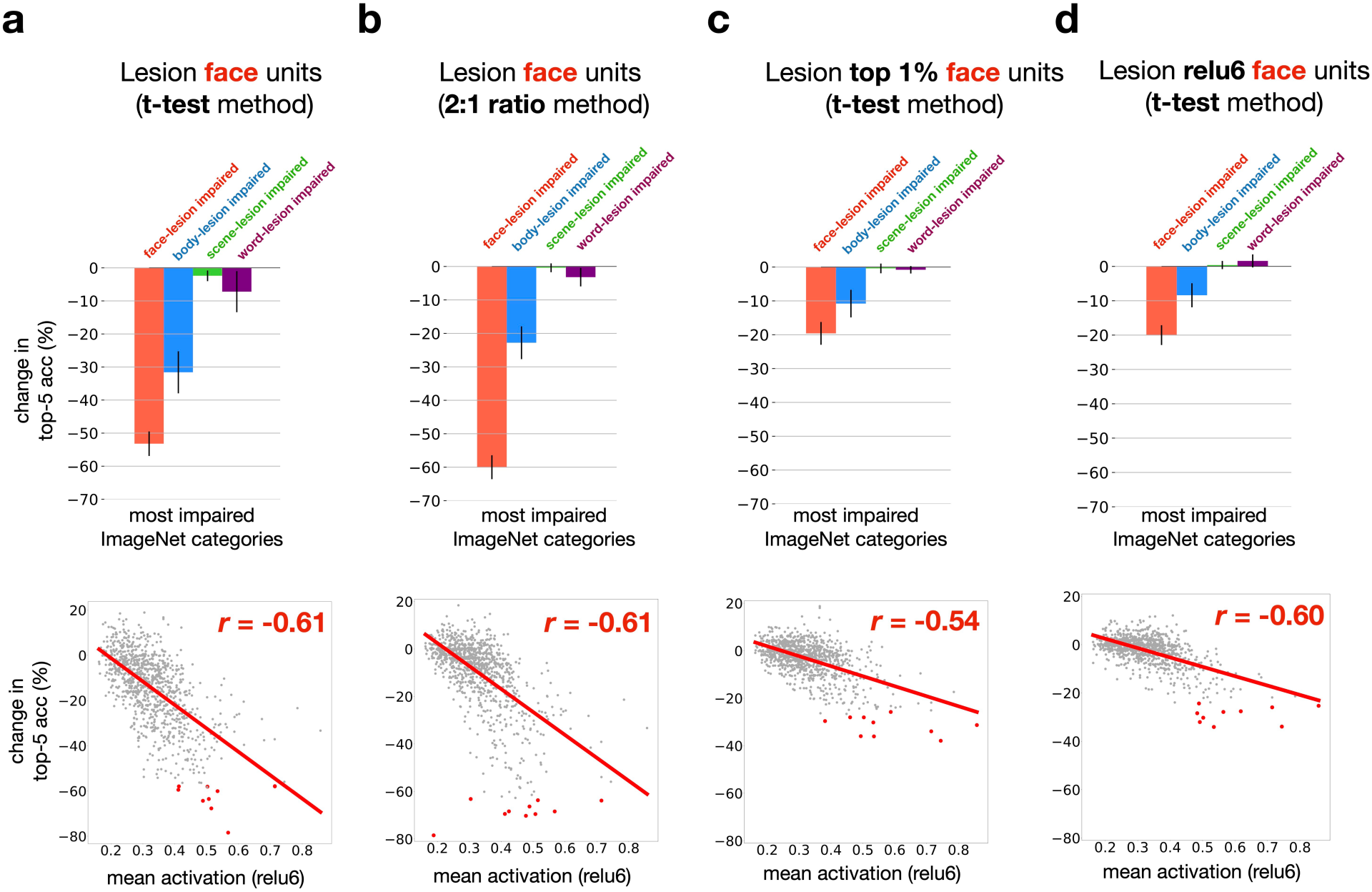
Testing alternative lesioning schemes and results from 2:1 selectivity criterion. (A) Top: changes in top-5 (%) accuracy are reported for the top 10 categories most impacted by face-selective unit lesions. Bottom: scatter plots show the relationship between mean activation pre-lesion and change in accuracy post-lesion for each of the 1000 ImageNet categories (dots). Same data as Figure 2C. Y values are jittered (values drawn from normal distribution; mean 0, std 0.5%) to enhance visibility of the results. (B) Results for face-selective units identified using the 2:1 ratio criterion, rather than the t-test + FDR localizer method. (C) Results for the t-test + FDR localizer method, with only the top 1% of most face-selective units in each relu layer lesioned. (D) Results for the t-test + FDR localizer method, with the full set of face-selective units from only layer relu6 lesioned.

**Supplementary Figure 8:**
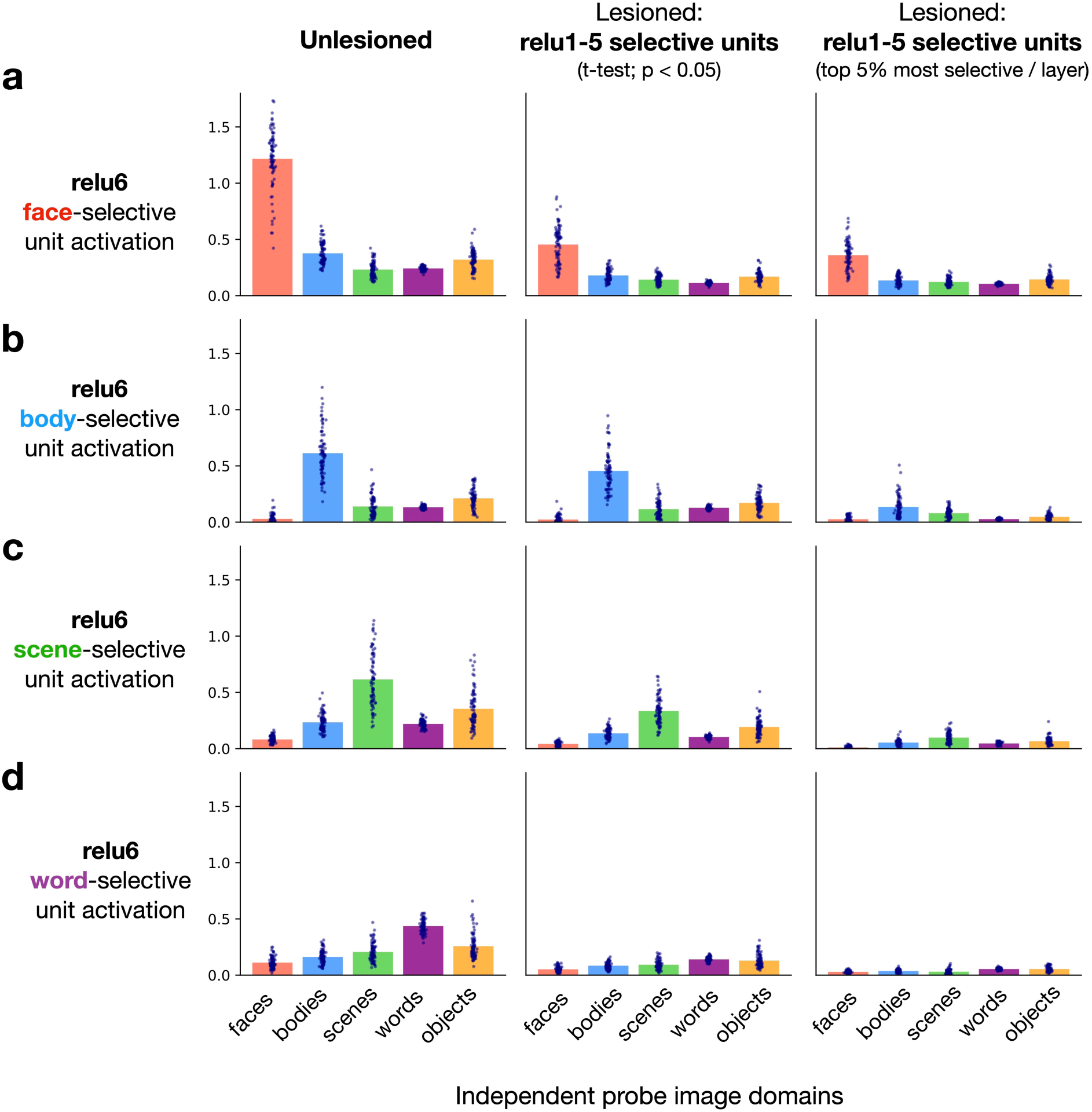
Impact of lesioning on downstream selectivity. (A) Mean activations are plotted in response to the RGB probe localizer images for face-selective units in layer relu6, identified using the t-test + FDR selectivity criterion. Left panel: activations with no lesions implemented (same data as **Supplementary Fig. 3B**). Middle panel: relu6 activations after lesions are applied to the face-selective units of relu layers 1-5, with layer relu6 unlesioned. Right panel: Same as middle panel, except with only the top 5% of selective units lesioned in each relu layer 1-5. (B-D) The same analyses, repeated for the body-, scene-, and word-selective units.

**Supplementary Figure 9:**
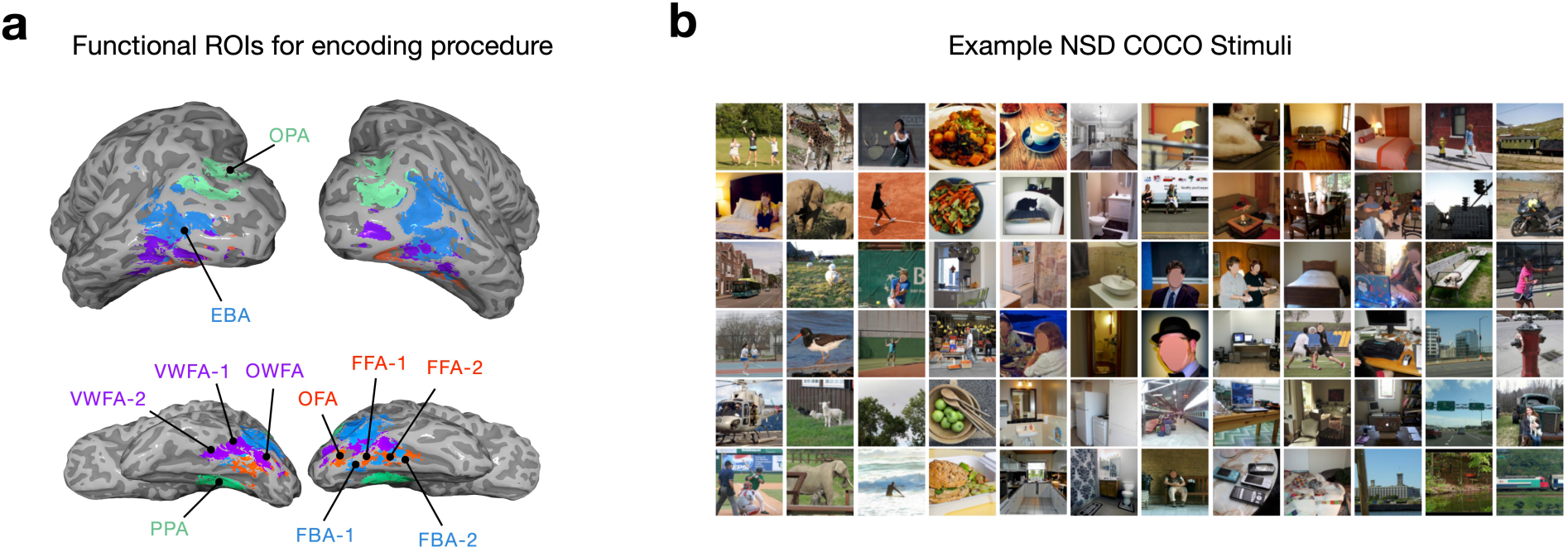
Overview of functional ROIs and NSD stimuli. (A) Data from the Natural Scenes Dataset (N = 8; representative subject plotted) used for the encoding procedure. 11 category selective ROIs are analyzed per subject, with preferences for faces (red), scenes (green), bodies (blue), and words (purple). (B) Examples from the test set of 515 stimuli from the NSD experiment. Photo credit: images in Panel B are samples from the public Microsoft COCO dataset, and those containing human faces are masked.

**Supplementary Figure 10:**
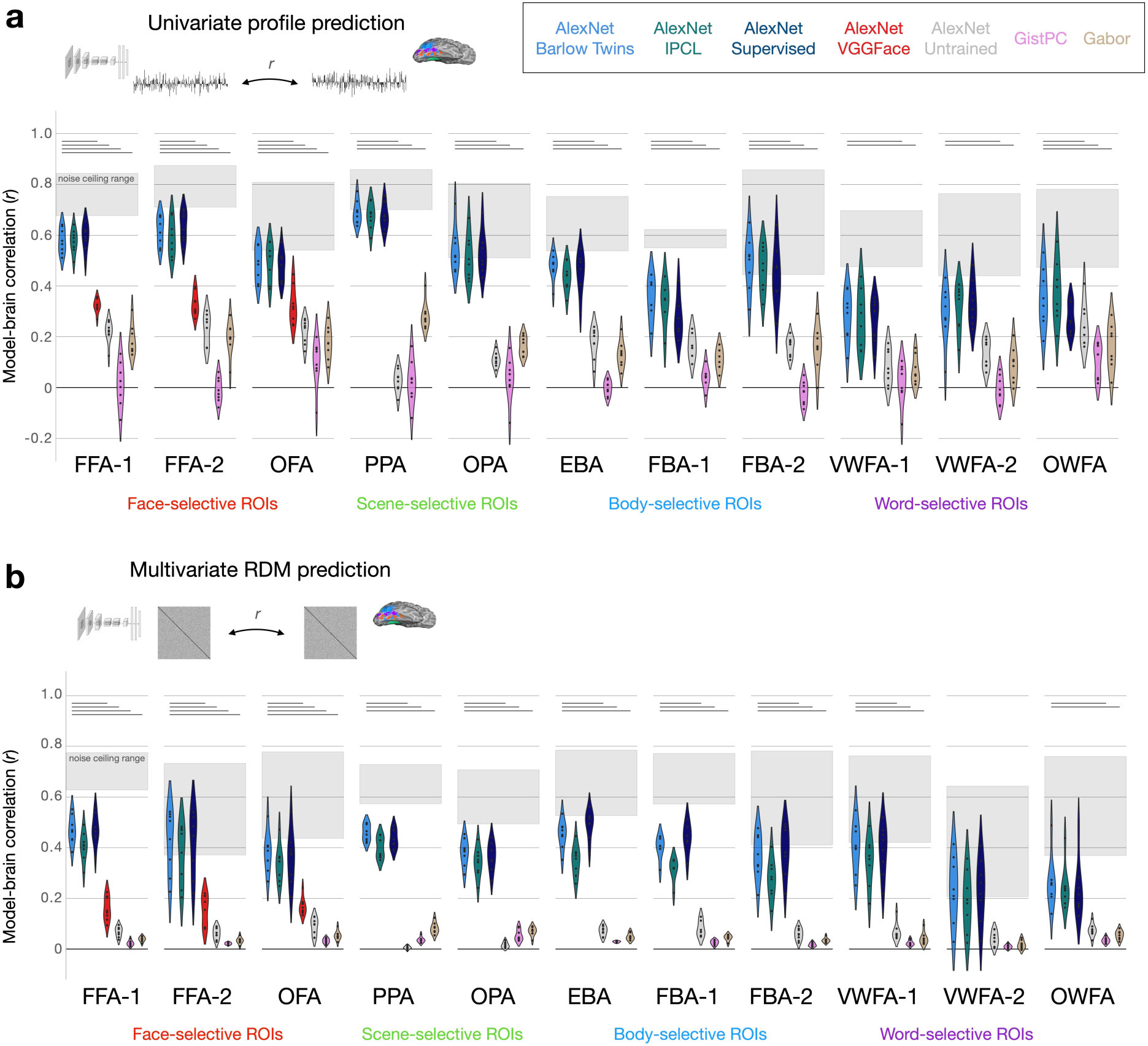
Brain prediction summary across a larger set of selective regions. Encoding results for predicting univariate response profiles (A) and multivariate RDMs (B) are summarized for the 8 NSD subjects, for each DNN and low-level image statistic model. Plotted values reflect best-layer correlations, as defined using cross-validation (see Methods) Shaded regions show the range of subject-specific noise ceilings computed over the data from each ROI. Significance is assessed using paired t-tests over subject-wise prediction levels between the AlexNet Barlow Twins model and each other candidate model. Horizontal bars reflect significant effects favoring the Barlow Twins model; Bonferroni-corrected p< 0.001.

**Supplementary Figure 11:**
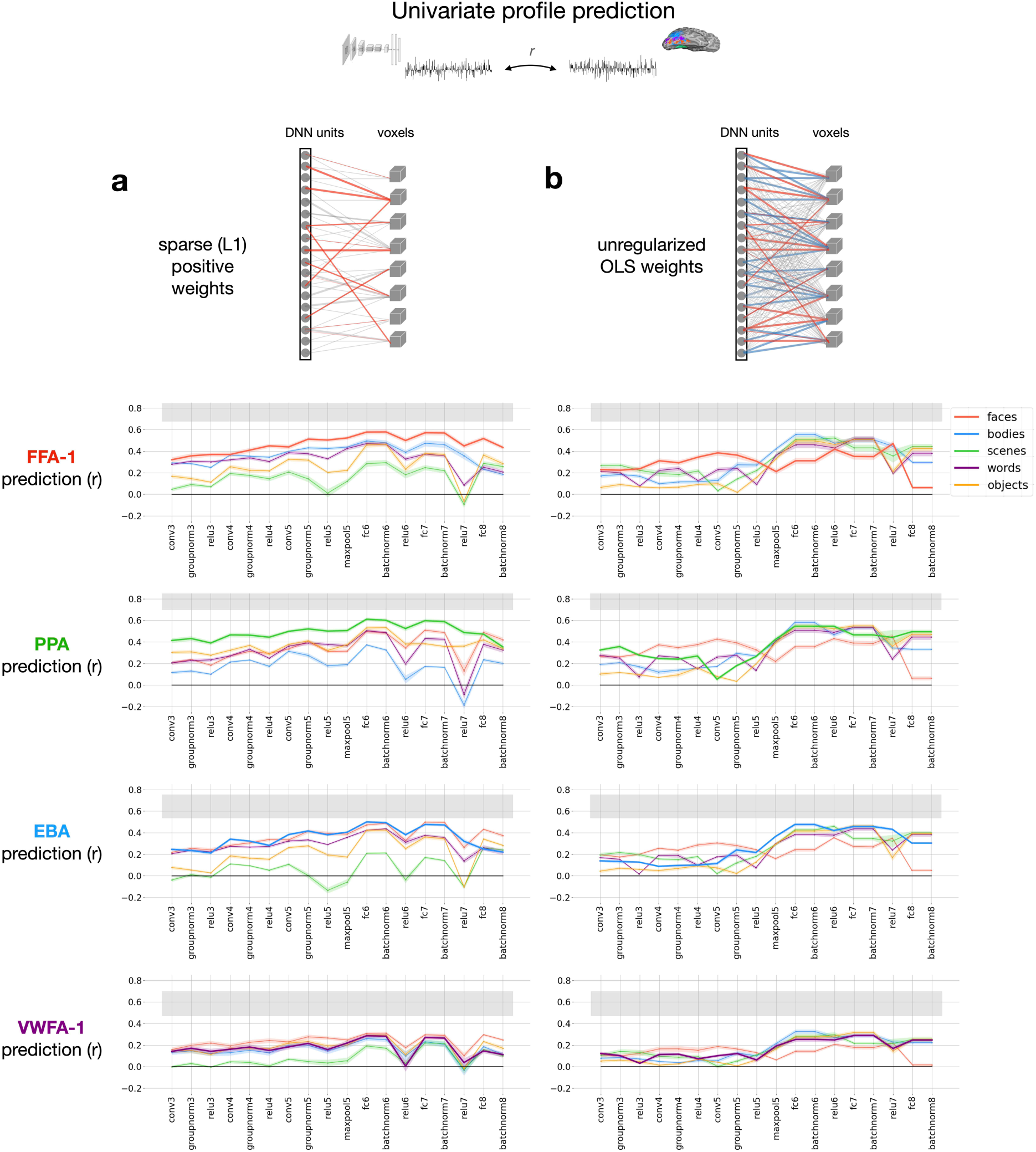
Impact of sparse-positive readout (univariate profile prediction). Layer summaries of encoding results are plotted for prediction of univariate response profiles in FFA-1, PPA, EBA, and VWFA-1 (N = 8 subjects, n = 515 test images). All data are from the AlexNet Barlow Twins model, with line colors representing the domain of selective units that are mapped onto the target ROI. Prediction results are compared between (A) the constrained encoding procedure involving sparse-positive weights only, and (B) an unregularized encoding procedure that uses ordinary least-squares (OLS) fitting, with no additional constraints. Shaded regions reflect the range of subject-specific noise ceilings computed over the univariate response profiles from each ROI.

**Supplementary Figure 12:**
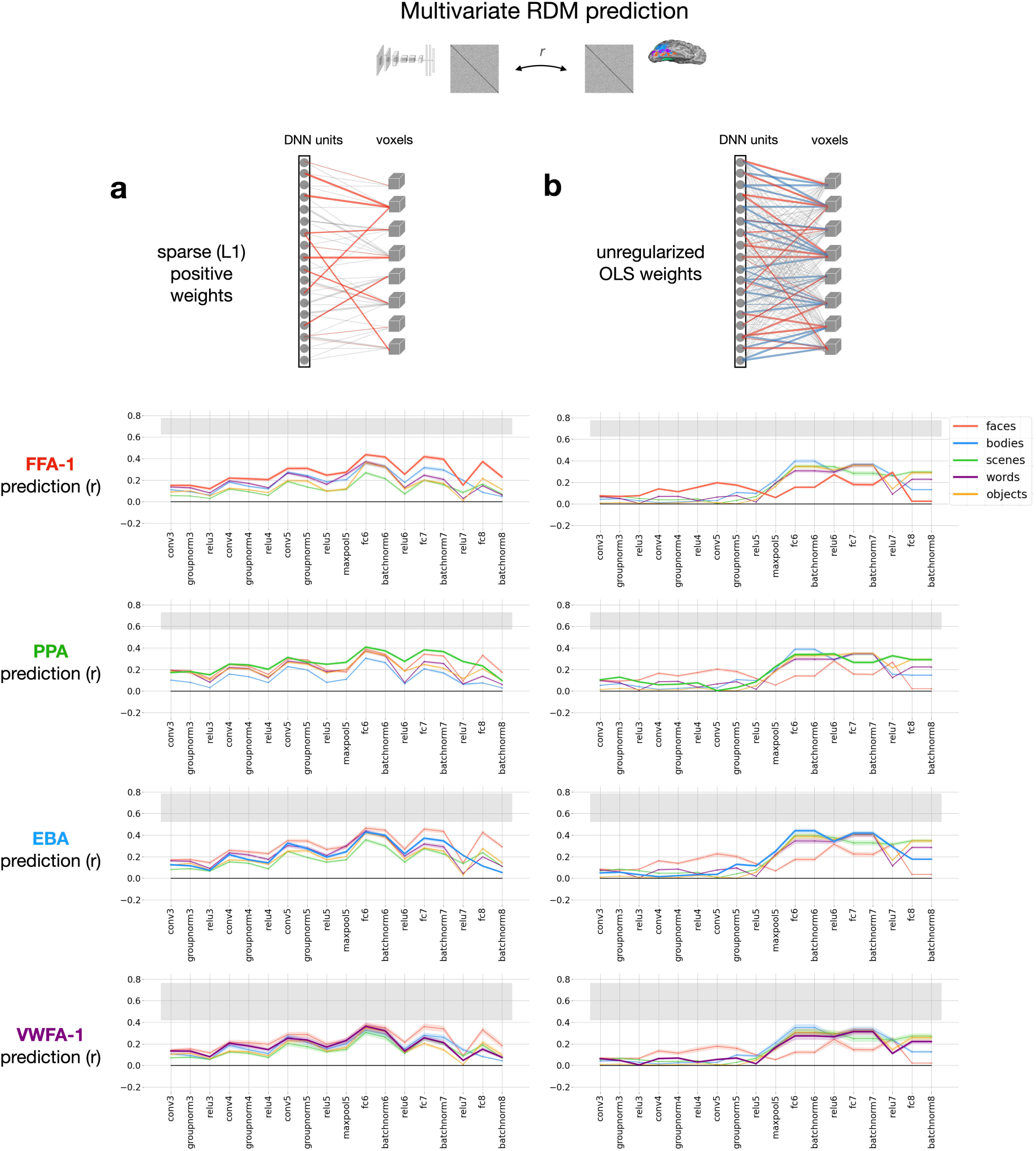
Impact of sparse-positive readout (multivariate RDM prediction). Layer summaries of encoding results are plotted for prediction of multivariate RDMs in FFA-1, PPA, EBA, and VWFA-1 (N = 8 subjects, n = 132,355 unique pairwise comparisons per RDM). All data are from the AlexNet Barlow Twins model, with line colors representing the domain of selective units that are mapped onto the target ROI. Prediction results are compared between (A) the constrained encoding procedure involving sparse-positive weights only, and (B) an unregularized encoding procedure that uses ordinary least-squares (OLS) fitting, with no additional constraints. Shaded regions reflect the range of subject-specific noise ceilings computed over the RDMs from each ROI.

**Supplementary Figure 13:**
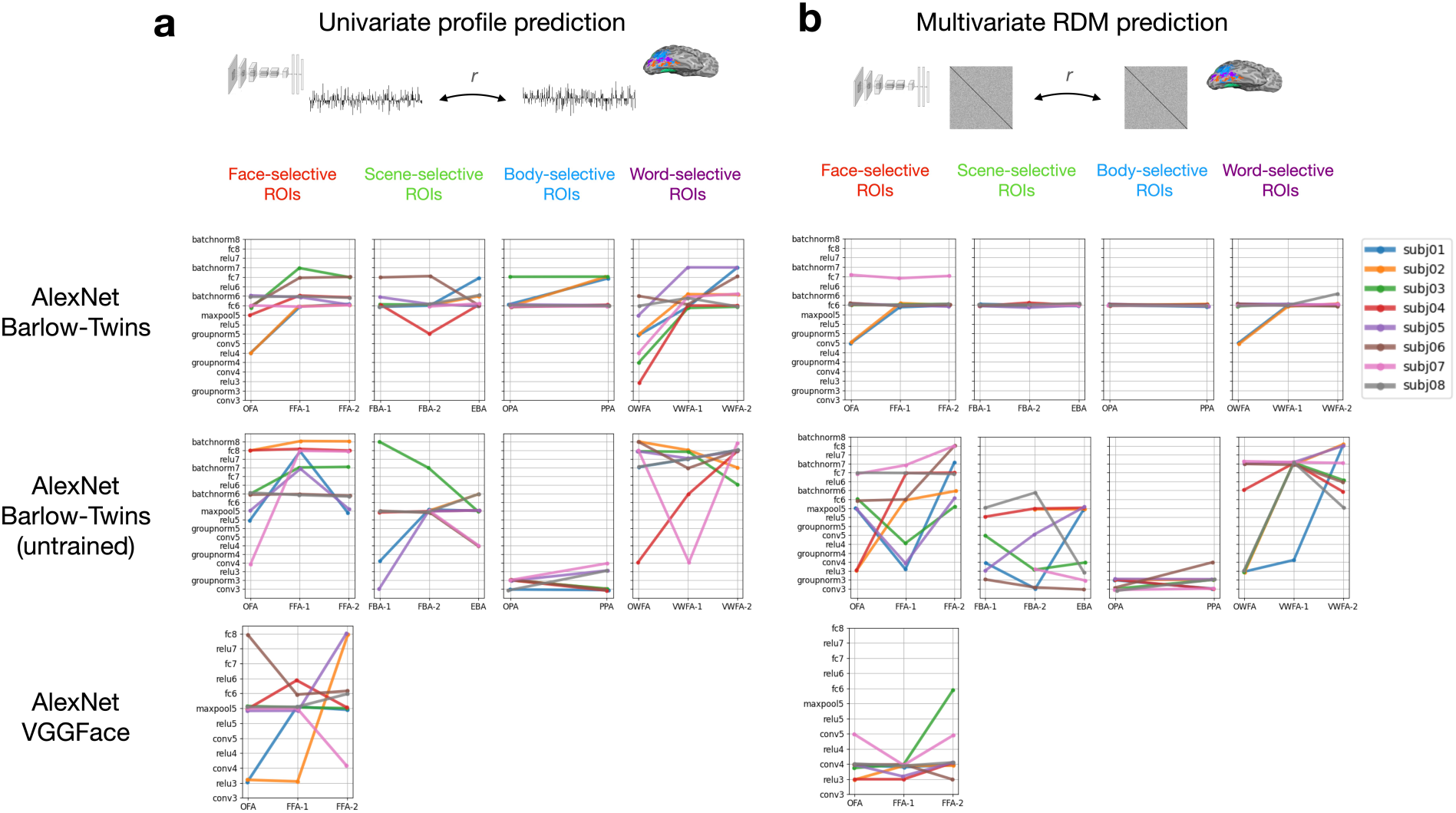
Indices of the most brain-predictive DNN layers. Indices of the most predictive DNN layers for univariate (A) and multivariate (B) encoding are identified using a validation set of 1000 subject-specific NSD stimuli. Lines reflect the outcomes for the 8 NSD subjects, across all 11 functional ROIs, for the three DNN models included in the encoding procedure.

## References

[1] M. Mishkin, L. G. Ungerleider, and K. A. Macko. Object vision and spatial vision: two cortical pathways. Trends in neurosciences, 6:414–417, 1983.

[2] J. V. Haxby, M. I. Gobbini, M. L. Furey, A. Ishai, J. L. Schouten, and P. Pietrini. Distributed and overlapping representations of faces and objects in ventral temporal cortex. Science, 293(5539):2425–2430, 2001.

[3] N. Kanwisher. Functional specificity in the human brain: a window into the functional architecture of the mind. Proceedings of the national academy of sciences, 107(25):11163–11170, 2010.

[4] K. Grill-Spector and K. S. Weiner. The functional architecture of the ventral temporal cortex and its role in categorization. Nature Reviews Neuroscience, 15(8):536–548, 2014.

[5] J. J. DiCarlo and D. D. Cox. Untangling invariant object recognition. Trends in cognitive sciences, 11(8):333–341, 2007.

[6] T. Meyer and N. C. Rust. Single-exposure visual memory judgments are reflected in inferotemporal cortex. elife, 7:e32259, 2018.

[7] N. Kanwisher, J. McDermott, and M. M. Chun. The fusiform face area: a module in human extrastriate cortex specialized for face perception. Journal of neuroscience, 17(11):4302–4311, 1997.

[8] G. McCarthy, A. Puce, J. C. Gore, and T. Allison. Face-specific processing in the human fusiform gyrus. Journal of cognitive neuroscience, 9(5):605–610, 1997.

[9] A. Puce, T. Allison, M. Asgari, J. C. Gore, and G. McCarthy. Differential sensitivity of human visual cortex to faces, letterstrings, and textures: a functional magnetic resonance imaging study. Journal of neuroscience, 16(16):5205–5215, 1996.

[10] D. Y. Tsao, W. A. Freiwald, R. B. Tootell, and M. S. Livingstone. A cortical region consisting entirely of face-selective cells. Science, 311(5761):670–674, 2006.

[11] P. E. Downing, Y. Jiang, M. Shuman, and N. Kanwisher. A cortical area selective for visual processing of the human body. Science, 293(5539):2470–2473, 2001.

[12] M. V. Peelen and P. E. Downing. Selectivity for the human body in the fusiform gyrus. Journal of neurophysiology, 93(1):603–608, 2005.

[13] R. Epstein and N. Kanwisher. A cortical representation of the local visual environment. Nature, 392(6676):598–601, 1998.

[14] G. K. Aguirre, E. Zarahn, and M. D’Esposito. An area within human ventral cortex sensitive to “building” stimuli: evidence and implications. Neuron, 21(2):373–383, 1998.

[15] S. Nasr, N. Liu, K. J. Devaney, X. Yue, R. Rajimehr, L. G. Ungerleider, and R. B. Tootell. Scene-selective cortical regions in human and nonhuman primates. Journal of Neuroscience, 31(39):13771–13785, 2011.

[16] S. E. Petersen, P. T. Fox, A. Z. Snyder, and M. E. Raichle. Activation of extrastriate and frontal cortical areas by visual words and word-like stimuli. Science, 249(4972):1041–1044, 1990.

[17] L. Cohen, S. Dehaene, L. Naccache, S. Lehéricy, G. Dehaene-Lambertz, M.-A. Hénaff, and F. Michel. The visual word form area: spatial and temporal characterization of an initial stage of reading in normal subjects and posterior split-brain patients. Brain, 123(2):291–307, 2000.

[18] T. A. Polk, M. Stallcup, G. K. Aguirre, D. C. Alsop, M. D’esposito, J. A. Detre, and M. J. Farah. Neural specialization for letter recognition. Journal of Cognitive Neuroscience, 14(2):145–159, 2002.

[19] H. P. Op de Beeck, I. Pillet, and J. B. Ritchie. Factors determining where category-selective areas emerge in visual cortex. Trends in cognitive sciences, 23(9):784–797, 2019.

[20] L. J. Powell, H. L. Kosakowski, and R. Saxe. Social origins of cortical face areas. Trends in cognitive sciences, 22(9):752–763, 2018.

[21] M. S. Livingstone, M. J. Arcaro, and P. F. Schade. Cortex is cortex: Ubiquitous principles drive face-domain development. Trends in cognitive sciences, 23(1):3, 2019.

[22] M. J. Arcaro and M. S. Livingstone. On the relationship between maps and domains in inferotemporal cortex. Nature Reviews Neuroscience, 22(9):573–583, 2021.

[23] R. A. Cowell and G. W. Cottrell. What evidence supports special processing for faces? a cautionary tale for fmri interpretation. Journal of Cognitive Neuroscience, 25(11):1777–1793, 2013.

[24] M. J. Tarr and I. Gauthier. Ffa: a flexible fusiform area for subordinate-level visual processing automatized by expertise. Nature neuroscience, 3(8):764–769, 2000.

[25] D. C. Plaut and M. Behrmann. Complementary neural representations for faces and words: A computational exploration. Cognitive neuropsychology, 28(3-4):251–275, 2011.

[26] S. He, H. Liu, Y. Jiang, C. Chen, Q. Gong, and X. Weng. Transforming a left lateral fusiform region into vwfa through training in illiterate adults. Journal of Vision, 9(8):853–853, 2009.

[27] T. Konkle and A. Oliva. A real-world size organization of object responses in occipitotemporal cortex. Neuron, 74(6):1114–1124, 2012.

[28] T. Konkle and A. Caramazza. The large-scale organization of object-responsive cortex is reflected in resting-state network architecture. Cerebral cortex, 27(10):4933–4945, 2017.

[29] N. A. Ratan Murty, P. Bashivan, A. Abate, J. J. DiCarlo, and N. Kanwisher. Computational models of category-selective brain regions enable high-throughput tests of selectivity. Nature communications, 12(1):5540, 2021.

[30] M. Khosla and L. Wehbe. High-level visual areas act like domain-general filters with strong selectivity and functional specialization. bioRxiv, pages 2022–03, 2022.

[31] H. L. Kosakowski, M. A. Cohen, A. Takahashi, B. Keil, N. Kanwisher, and R. Saxe. Selective responses to faces, scenes, and bodies in the ventral visual pathway of infants. Current Biology, 32(2):265–274, 2022.

[32] N. Kanwisher. Domain specificity in face perception. Nature neuroscience, 3(8):759–763, 2000.

[33] B. Z. Mahon and A. Caramazza. What drives the organization of object knowledge in the brain? Trends in cognitive sciences, 15(3):97–103, 2011.

[34] M. V. Peelen and P. E. Downing. Category selectivity in human visual cortex: Beyond visual object recognition. Neuropsychologia, 105:177–183, 2017.

[35] S. Bracci, J. B. Ritchie, and H. O. de Beeck. On the partnership between neural representations of object categories and visual features in the ventral visual pathway. Neuropsychologia, 105:153– 164, 2017.

[36] S. Moeller, W. A. Freiwald, and D. Y. Tsao. Patches with links: a unified system for processing faces in the macaque temporal lobe. Science, 320(5881):1355–1359, 2008.

[37] O. Pascalis and D. J. Kelly. The origins of face processing in humans: Phylogeny and ontogeny. Perspectives on psychological science, 4(2):200–209, 2009.

[38] F. S. Kamps, C. L. Hendrix, P. A. Brennan, and D. D. Dilks. Connectivity at the origins of domain specificity in the cortical face and place networks. Proceedings of the National Academy of Sciences, 117(11):6163–6169, 2020.

[39] B. Z. Mahon. Domain-specific connectivity drives the organization of object knowledge in the brain. In Handbook of Clinical Neurology, volume 187, pages 221–244. Elsevier, 2022.

[40] T. Hannagan, A. Amedi, L. Cohen, G. Dehaene-Lambertz, and S. Dehaene. Origins of the specialization for letters and numbers in ventral occipitotemporal cortex. Trends in cognitive sciences, 19(7):374–382, 2015.

[41] S. Dehaene and G. Dehaene-Lambertz. Is the brain prewired for letters? Nature neuroscience, 19(9):1192–1193, 2016.

[42] J. Li, D. E. Osher, H. A. Hansen, and Z. M. Saygin. Innate connectivity patterns drive the development of the visual word form area. Scientific reports, 10(1):18039, 2020.

[43] M. Habib and A. Sirigu. Pure topographical disorientation: a definition and anatomical basis. Cortex, 23(1):73–85, 1987.

[44] R. Epstein, E. A. DeYoe, D. Z. Press, A. C. Rosen, and N. Kanwisher. Neuropsychological evidence for a topographical learning mechanism in parahippocampal cortex. Cognitive neuropsychology, 18(6):481–508, 2001.

[45] J. J. Barton, D. Z. Press, J. P. Keenan, and M. O’Connor. Lesions of the fusiform face area impair perception of facial configuration in prosopagnosia. Neurology, 58(1):71–78, 2002.

[46] C. Schiltz, B. Sorger, R. Caldara, F. Ahmed, E. Mayer, R. Goebel, and B. Rossion. Impaired face discrimination in acquired prosopagnosia is associated with abnormal response to individual faces in the right middle fusiform gyrus. Cerebral Cortex, 16(4):574–586, 2006.

[47] N. Lang, J. Baudewig, K. Kallenberg, A. Antal, S. Happe, P. Dechent, and W. Paulus. Transient prosopagnosia after ischemic stroke. Neurology, 66(6):916–916, 2006.

[48] J. J. Barton. Structure and function in acquired prosopagnosia: lessons from a series of 10 patients with brain damage. Journal of neuropsychology, 2(1):197–225, 2008.

[49] L. Dricot, B. Sorger, C. Schiltz, R. Goebel, and B. Rossion. The roles of “face” and “nonface” areas during individual face perception: evidence by fmri adaptation in a brain-damaged prosopagnosic patient. Neuroimage, 40(1):318–332, 2008.

[50] V. Moro, C. Urgesi, S. Pernigo, P. Lanteri, M. Pazzaglia, and S. M. Aglioti. The neural basis of body form and body action agnosia. Neuron, 60(2):235–246, 2008.

[51] C. Urgesi, G. Berlucchi, and S. M. Aglioti. Magnetic stimulation of extrastriate body area impairs visual processing of nonfacial body parts. Current Biology, 14(23):2130–2134, 2004.

[52] D. Pitcher, V. Walsh, G. Yovel, and B. Duchaine. Tms evidence for the involvement of the right occipital face area in early face processing. Current Biology, 17(18):1568–1573, 2007.

[53] D. Pitcher, L. Charles, J. T. Devlin, V. Walsh, and B. Duchaine. Triple dissociation of faces, bodies, and objects in extrastriate cortex. Current Biology, 19(4):319–324, 2009.

[54] S. Moeller, T. Crapse, L. Chang, and D. Y. Tsao. The effect of face patch microstimulation on perception of faces and objects. Nature neuroscience, 20(5):743–752, 2017.

[55] S. Sadagopan, W. Zarco, and W. A. Freiwald. A causal relationship between face-patch activity and face-detection behavior. Elife, 6:e18558, 2017.

[56] N. Kanwisher and J. J. Barton. The functional architecture of the face system: Integrating evidence from fmri and patient studies. The Oxford handbook of face perception, pages 111–129, 2011.

[57] C. G. Gross. Genealogy of the “grandmother cell”. The Neuroscientist, 8(5):512–518, 2002.

[58] A. Ishai, L. G. Ungerleider, A. Martin, J. L. Schouten, and J. V. Haxby. Distributed representation of objects in the human ventral visual pathway. Proceedings of the National Academy of Sciences, 96(16):9379–9384, 1999.

[59] P. E. Downing, A.-Y. Chan, M. V. Peelen, C. Dodds, and N. Kanwisher. Domain specificity in visual cortex. Cerebral cortex, 16(10):1453–1461, 2006.

[60] R. Kiani, H. Esteky, K. Mirpour, and K. Tanaka. Object category structure in response patterns of neuronal population in monkey inferior temporal cortex. Journal of neurophysiology, 97(6):4296– 4309, 2007.

[61] A. Ishai, L. G. Ungerleider, A. Martin, and J. V. Haxby. The representation of objects in the human occipital and temporal cortex. Journal of Cognitive Neuroscience, 12(Supplement 2):35–51, 2000.

[62] N. Kriegeskorte, M. Mur, D. A. Ruff, R. Kiani, J. Bodurka, H. Esteky, K. Tanaka, and P. A. Bandettini. Matching categorical object representations in inferior temporal cortex of man and monkey. Neuron, 60(6):1126–1141, 2008.

[63] M. Mur, D. A. Ruff, J. Bodurka, P. De Weerd, P. A. Bandettini, and N. Kriegeskorte. Categorical, yet graded–single-image activation profiles of human category-selective cortical regions. Journal of Neuroscience, 32(25):8649–8662, 2012.

[64] I. Biederman. Recognition-by-components: a theory of human image understanding. Psychological review, 94(2):115, 1987.

[65] S. Chung and L. Abbott. Neural population geometry: An approach for understanding biological and artificial neural networks. Current opinion in neurobiology, 70:137–144, 2021.

[66] D. D. Cox and R. L. Savoy. Functional magnetic resonance imaging (fmri)“brain reading”: detecting and classifying distributed patterns of fmri activity in human visual cortex. Neuroimage, 19(2):261–270, 2003.

[67] E. Eger, J. Ashburner, J.-D. Haynes, R. J. Dolan, and G. Rees. fmri activity patterns in human loc carry information about object exemplars within category. Journal of cognitive neuroscience, 20(2):356–370, 2008.

[68] A. C. Connolly, J. S. Guntupalli, J. Gors, M. Hanke, Y. O. Halchenko, Y.-C. Wu, H. Abdi, and J. V. Haxby. The representation of biological classes in the human brain. Journal of Neuroscience, 32(8):2608–2618, 2012.

[69] R. F. Schwarzlose, J. D. Swisher, S. Dang, and N. Kanwisher. The distribution of category and location information across object-selective regions in human visual cortex. Proceedings of the National Academy of Sciences, 105(11):4447–4452, 2008.

[70] M. Spiridon and N. Kanwisher. How distributed is visual category information in human occipito-temporal cortex? an fmri study. Neuron, 35(6):1157–1165, 2002.

[71] M. A. Williams, S. Dang, and N. G. Kanwisher. Only some spatial patterns of fmri response are read out in task performance. Nature neuroscience, 10(6):685–686, 2007.

[72] L. Reddy and N. Kanwisher. Category selectivity in the ventral visual pathway confers robustness to clutter and diverted attention. Current Biology, 17(23):2067–2072, 2007.

[73] G. Schalk, C. Kapeller, C. Guger, H. Ogawa, S. Hiroshima, R. Lafer-Sousa, Z. M. Saygin, K. Kamada, and N. Kanwisher. Facephenes and rainbows: Causal evidence for functional and anatomical specificity of face and color processing in the human brain. Proceedings of the National Academy of Sciences, 114(46):12285–12290, 2017.

[74] P. Bao, L. She, M. McGill, and D. Y. Tsao. A map of object space in primate inferotemporal cortex. Nature, 583(7814):103–108, 2020.

[75] F. R. Doshi and T. Konkle. Cortical topographic motifs emerge in a self-organized map of object space. Science Advances, 9(25):eade8187, 2023.

[76] E. Margalit, H. Lee, D. Finzi, J. J. DiCarlo, K. Grill-Spector, and D. L. Yamins. A unifying principle for the functional organization of visual cortex. bioRxiv, pages 2023–05, 2023.

[77] J. Zbontar, L. Jing, I. Misra, Y. LeCun, and S. Deny. Barlow twins: Self-supervised learning via redundancy reduction. arXiv preprint arXiv:2103.03230, 2021.

[78] T. Konkle and G. A. Alvarez. A self-supervised domain-general learning framework for human ventral stream representation. Nature communications, 13(1):491, 2022.

[79] A. Krizhevsky, I. Sutskever, and G. E. Hinton. ImageNet classification with deep convolutional neural networks. In Advances in Neural Information Processing Systems, pages 1097–1105, 2012.

80. J. Deng, W. Dong, R. Socher, L.-J. Li, K. Li, and L. Fei-Fei. Imagenet: A large-scale hierarchical image database. In 2009 IEEE conference on computer vision and pattern recognition, pages 248–255. Ieee, 2009.

[81] Z. Wu, Y. Xiong, S. X. Yu, and D. Lin. Unsupervised feature learning via non-parametric instance discrimination. In Proceedings of the IEEE conference on computer vision and pattern recognition, pages 3733–3742, 2018.

82. T. Chen, S. Kornblith, M. Norouzi, and G. Hinton. A simple framework for contrastive learning of visual representations. In International conference on machine learning, pages 1597–1607. PMLR, 2020.

[83] M. Caron, I. Misra, J. Mairal, P. Goyal, P. Bojanowski, and A. Joulin. Unsupervised learning of visual features by contrasting cluster assignments. NeurIPS, 2020. https://arxiv.org/abs/2006.09882.

[84] J.-B. Grill, F. Strub, F. Altché, C. Tallec, P. H. Richemond, E. Buchatskaya, C. Doersch, B. A. Pires, Z. D. Guo, M. G. Azar, et al. Bootstrap your own latent: A new approach to self-supervised learning. arXiv preprint arXiv:2006.07733, 2020.

[85] Y. Shu, X. Gu, G.-Z. Yang, and B. Lo. Revisiting self-supervised contrastive learning for facial expression recognition. arXiv preprint arXiv:2210.03853, 2022.

[86] H. Wang, V. Sanchez, and C.-T. Li. Cross-age contrastive learning for age-invariant face recognition. In ICASSP 2024-2024 IEEE International Conference on Acoustics, Speech and Signal Processing (ICASSP), pages 4600–4604. IEEE, 2024.

[87] Q. Garrido, Y. Chen, A. Bardes, L. Najman, and Y. Lecun. On the duality between contrastive and non-contrastive self-supervised learning. arXiv preprint arXiv:2206.02574, 2022.

[88] C. Tao, H. Wang, X. Zhu, J. Dong, S. Song, G. Huang, and J. Dai. Exploring the equivalence of siamese self-supervised learning via a unified gradient framework. In Proceedings of the IEEE/CVF Conference on Computer Vision and Pattern Recognition, pages 14431–14440, 2022.

[89] W. Huang, M. Yi, and X. Zhao. Towards the generalization of contrastive self-supervised learning. arXiv preprint arXiv:2111.00743, 2021.

[90] A. Stigliani, K. S. Weiner, and K. Grill-Spector. Temporal processing capacity in high-level visual cortex is domain specific. Journal of Neuroscience, 35(36):12412–12424, 2015.

[91] E. J. Allen, G. St-Yves, Y. Wu, J. L. Breedlove, J. S. Prince, L. T. Dowdle, M. Nau, B. Caron, F. Pestilli, I. Charest, et al. A massive 7t fmri dataset to bridge cognitive neuroscience and artificial intelligence. Nature neuroscience, 25(1):116–126, 2022.

[92] K. Grill-Spector, N. Knouf, and N. Kanwisher. The fusiform face area subserves face perception, not generic within-category identification. Nature neuroscience, 7(5):555–562, 2004.

[93] R. Geirhos, K. Narayanappa, B. Mitzkus, M. Bethge, F. A. Wichmann, and W. Brendel. On the surprising similarities between supervised and self-supervised models. arXiv preprint arXiv:2010.08377, 2020.

[94] S. Baek, M. Song, J. Jang, G. Kim, and S.-B. Paik. Face detection in untrained deep neural networks. Nature communications, 12(1):7328, 2021.

[95] T. Konkle and A. Caramazza. Tripartite organization of the ventral stream by animacy and object size. Journal of Neuroscience, 33(25):10235–10242, 2013.

[96] B. Zhang, S. He, and X. Weng. Localization and functional characterization of an occipital visual word form sensitive area. Scientific reports, 8(1):6723, 2018.

[97] L. Chang and D. Y. Tsao. The code for facial identity in the primate brain. Cell, 169(6):1013– 1028, 2017.

[98] I. Yildirim, M. Belledonne, W. Freiwald, and J. Tenenbaum. Efficient inverse graphics in biological face processing. Science advances, 6(10):eaax5979, 2020.

[99] I. Higgins, L. Chang, V. Langston, D. Hassabis, C. Summerfield, D. Tsao, and M. Botvinick. Unsupervised deep learning identifies semantic disentanglement in single inferotemporal face patch neurons. Nature communications, 12(1):6456, 2021.

[100] G. J. Edwards, T. F. Cootes, and C. J. Taylor. Face recognition using active appearance models. In Computer Vision—ECCV’98: 5th European Conference on Computer Vision Freiburg, Germany, June 2–6, 1998 Proceedings*, Volume* II *5*, pages 581–595. Springer, 1998.

[101] J. S. Prince, C. Conwell, G. A. Alvarez, and T. Konkle. A case for sparse positive alignment of neural systems. In ICLR 2024 Workshop on Representational Alignment, 2024.

102. Q. Cao, L. Shen, W. Xie, O. M. Parkhi, and A. Zisserman. Vggface2: A dataset for recognising faces across pose and age. In 2018 13th IEEE international conference on automatic face & gesture recognition (FG 2018), pages 67–74. IEEE, 2018.

[103] C. Conwell, J. S. Prince, K. N. Kay, G. A. Alvarez, and T. Konkle. What can 1.8 billion regressions tell us about the pressures shaping high-level visual representation in brains and machines? bioRxiv, pages 2022–03, 2022.

[104] K. Vinken, J. S. Prince, T. Konkle, and M. Livingstone. The neural code for ‘face cells’ is not face specific. bioRxiv, pages 2022–03, 2022.

[105] M. A. Cohen, G. A. Alvarez, K. Nakayama, and T. Konkle. Visual search for object categories is predicted by the representational architecture of high-level visual cortex. Journal of neurophysiology, 117(1):388–402, 2017.

[106] C. P. Hung, G. Kreiman, T. Poggio, and J. J. DiCarlo. Fast readout of object identity from macaque inferior temporal cortex. Science, 310(5749):863–866, 2005.

[107] T. Naselaris, K. N. Kay, S. Nishimoto, and J. L. Gallant. Encoding and decoding in fmri. Neuroimage, 56(2):400–410, 2011.

[108] A. Mahmoudi, S. Takerkart, F. Regragui, D. Boussaoud, A. Brovelli, et al. Multivoxel pattern analysis for fmri data: a review. Computational and mathematical methods in medicine, 2012, 2012.

[109] S. Bracci and M. V. Peelen. Body and object effectors: the organization of object representations in high-level visual cortex reflects body–object interactions. Journal of Neuroscience, 33(46):18247–18258, 2013.

[110] S. Bracci, A. Caramazza, and M. V. Peelen. Representational similarity of body parts in human occipitotemporal cortex. Journal of Neuroscience, 35(38):12977–12985, 2015.

[111] M. F. Wurm, A. Caramazza, and A. Lingnau. Action categories in lateral occipitotemporal cortex are organized along sociality and transitivity. Journal of Neuroscience, 37(3):562–575, 2017.

[112] L. Tarhan and T. Konkle. Sociality and interaction envelope organize visual action representations. Nature Communications, 11(1):3002, 2020.

[113] L. Tarhan, J. De Freitas, and T. Konkle. Behavioral and neural representations en route to intuitive action understanding. Neuropsychologia, 163:108048, 2021.

[114] E. Abassi and L. Papeo. The representation of two-body shapes in the human visual cortex. Journal of Neuroscience, 40(4):852–863, 2020.

[115] J. Mehrer, C. J. Spoerer, E. C. Jones, N. Kriegeskorte, and T. C. Kietzmann. An ecologically motivated image dataset for deep learning yields better models of human vision. Proceedings of the National Academy of Sciences, 118(8):e2011417118, 2021.

[116] J. Sullivan, M. Mei, A. Perfors, E. Wojcik, and M. C. Frank. Saycam: A large, longitudinal audiovisual dataset recorded from the infant’s perspective. Open mind, 5:20–29, 2021.

117. K. Yang, J. Yau, L. Fei-Fei, J. Deng, and O. Russakovsky. A study of face obfuscation in imagenet. In International Conference on Machine Learning (ICML).

[118] R. Le Grand, C. J. Mondloch, D. Maurer, and H. P. Brent. Early visual experience and face processing. Nature, 410(6831):890–890, 2001.

[119] L. B. Smith. Learning to recognize objects. Psychological Science, 14(3):244–250, 2003.

[120] G. Raz and R. Saxe. Learning in infancy is active, endogenously motivated, and depends on the prefrontal cortices. Annual Review of Developmental Psychology, 2:247–268, 2020.

[121] A. F. Pereira and L. B. Smith. Developmental changes in visual object recognition between 18 and 24 months of age. Developmental science, 12(1):67–80, 2009.

[122] Y. Ostrovsky, E. Meyers, S. Ganesh, U. Mathur, and P. Sinha. Visual parsing after recovery from blindness. Psychological Science, 20(12):1484–1491, 2009.

[123] E. S. Spelke. What Babies Know: Core Knowledge and Composition Volume 1, volume 1. Oxford University Press, 2022.

[124] K. Dobs, J. Yuan, J. Martinez, and N. Kanwisher. Behavioral signatures of face perception emerge in deep neural networks optimized for face recognition. Proceedings of the National Academy of Sciences, 120(32):e2220642120, 2023.

[125] N. M. Blauch, M. Behrmann, and D. C. Plaut. Computational insights into human perceptual expertise for familiar and unfamiliar face recognition. Cognition, 208:104341, 2021.

[126] L. Chang, B. Egger, T. Vetter, and D. Y. Tsao. Explaining face representation in the primate brain using different computational models. Current Biology, 31(13):2785–2795, 2021.

[127] S. Grossman, G. Gaziv, E. M. Yeagle, M. Harel, P. Mégevand, D. M. Groppe, S. Khuvis, J. L. Herrero, M. Irani, A. D. Mehta, et al. Convergent evolution of face spaces across human face-selective neuronal groups and deep convolutional networks. Nature communications, 10(1):4934, 2019.

[128] K. R. Storrs, T. C. Kietzmann, A. Walther, J. Mehrer, and N. Kriegeskorte. Diverse deep neural networks all predict human inferior temporal cortex well, after training and fitting. Journal of Cognitive Neuroscience, 33(10):2044–2064, 2021.

[129] U. Gü çlü and M. A. van Gerven. Deep neural networks reveal a gradient in the complexity of neural representations across the ventral stream. Journal of Neuroscience, 35(27):10005–10014, 2015.

[130] P. Agrawal, D. Stansbury, J. Malik, and J. L. Gallant. Pixels to voxels: Modeling visual representation in the human brain, 2014.

[131] M. Eickenberg, A. Gramfort, G. Varoquaux, and B. Thirion. Seeing it all: Convolutional network layers map the function of the human visual system. NeuroImage, 152:184–194, 2017.

[132] T. D. la Tour, M. Lu, M. Eickenberg, and J. L. Gallant. A finer mapping of convolutional neural network layers to the visual cortex. In SVRHM 2021 Workshop@ NeurIPS, 2021.

[133] A. H. Williams, E. Kunz, S. Kornblith, and S. Linderman. Generalized shape metrics on neural representations. Advances in Neural Information Processing Systems, 34:4738–4750, 2021.

[134] M. Khosla and A. H. Williams. Soft matching distance: A metric on neural representations that captures single-neuron tuning. arXiv preprint arXiv:2311.09466, 2023.

135. I. Sucholutsky, L. Muttenthaler, A. Weller, A. Peng, A. Bobu, B. Kim, B. C. Love, E. Grant, J. Achterberg, J. B. Tenenbaum, et al. Getting aligned on representational alignment. arXiv preprint arXiv:2310.13018, 2023.

[136] A. Jaegle, V. Mehrpour, Y. Mohsenzadeh, T. Meyer, A. Oliva, and N. Rust. Population response magnitude variation in inferotemporal cortex predicts image memorability. Elife, 8:e47596, 2019.

137. T.-Y. Lin, M. Maire, S. Belongie, J. Hays, P. Perona, D. Ramanan, P. Dollár, and C. L. Zitnick. Microsoft coco: Common objects in context. In European conference on computer vision, pages 740–755. Springer, 2014.

[138] J. S. Prince, I. Charest, J. W. Kurzawski, J. A. Pyles, M. J. Tarr, and K. N. Kay. Improving the accuracy of single-trial fmri response estimates using glmsingle. Elife, 11:e77599, 2022.

[139] L. Tarhan and T. Konkle. Reliability-based voxel selection. Neuroimage, 207:116350, 2020.

[140] A. Oliva and A. Torralba. Building the gist of a scene: The role of global image features in recognition. Progress in brain research, 155:23–36, 2006.

[141] K. Kay, J. S. Prince, T. Gebhart, G. Tuckute, J. Zhou, T. Naselaris, and H. Schutt. Disentangling signal and noise in neural responses through generative modeling. bioRxiv, pages 2024–04, 2024.

[142] A. Kallmayer, J. Prince, and T. Konkle. Comparing representations that support object, scene, and face recognition using representational trajectory analysis. Journal of Vision, 20(11):861–861, 2020.

